# PICRUSt2: An improved and customizable approach for metagenome inference

**DOI:** 10.1101/672295

**Authors:** Gavin M. Douglas, Vincent J. Maffei, Jesse Zaneveld, Svetlana N. Yurgel, James R. Brown, Christopher M. Taylor, Curtis Huttenhower, Morgan G. I. Langille

## Abstract

One major limitation of microbial community marker gene sequencing is that it does not provide direct information on the functional composition of sampled communities. Here, we present PICRUSt2 (https://github.com/picrust/picrust2), which expands the capabilities of the original PICRUSt method^1^ to predict the functional potential of a community based on marker gene sequencing profiles. This updated method and implementation includes several improvements over the previous algorithm: an expanded database of gene families and reference genomes, a new approach now compatible with any OTU-picking or denoising algorithm, and novel phenotype predictions. Upon evaluation, PICRUSt2 was more accurate than PICRUSt1 and other current approaches overall. PICRUSt2 is also now more flexible and allows the addition of custom reference databases. We highlight these improvements and also important caveats regarding the use of predicted metagenomes, which are related to the inherent challenges of analyzing metagenome data in general.

---

The most common approach for profiling communities is to sequence the highly conserved 16S rRNA gene. Functional profiles cannot be directly identified from 16S rRNA gene sequence data due to strain variation and because 16S rRNA genes are not unique among microbes, but several approaches have been developed to infer approximate microbial community functions from taxonomic profiles (and thus amplicon sequences) alone^1–6^. Importantly, these methods predict functional potential, i.e. functions encoded at the level of DNA. Although shotgun metagenomic sequencing (MGS) directly samples genetic functional potential within microbial communities, this methodology is not without limitations. In particular, functional inference from amplicon data remains important for samples with substantial host contamination (e.g. biopsy samples), low biomass, and where metagenomic sequencing is not economically feasible.

PICRUSt^1^ (hereafter “PICRUSt1”) was the first tool developed and the most widely used for metagenome prediction, but like any inference model has several limitations. First, the standard PICRUSt1 workflow requires input sequences to be operational taxonomic units (OTUs) generated from closed-reference OTU picking against a compatible version of the Greengenes database^7^. Due to this limitation, the default PICRUSt1 workflow is incompatible with sequence denoising methods^8^, which produce amplicon sequence variants (ASVs) rather than OTUs. ASVs have finer resolution, allowing closely related organisms to be more readily distinguished. Lastly, the prokaryotic reference databases used by PICRUSt1 have not been updated since 2013 and lack many recently added gene families and pathway mappings.

The PICRUSt2 algorithm includes new steps that optimize genome prediction, which we hypothesized would improve prediction accuracy (**Fig 1**). These are: (1) study sequences are now placed into a pre-existing phylogeny rather than relying on discrete predictions limited to reference OTUs (**Fig 1b**); (2) predictions are based off of a greatly increased number of reference genomes and gene families (**Fig 1c**); (3) pathway abundance inference is now more stringently performed (**Supp Fig 1**); (4) predictions can now be made for higher level phenotypes; and (5) custom databases are easier to integrate into the prediction pipeline.

**Figure 1:**
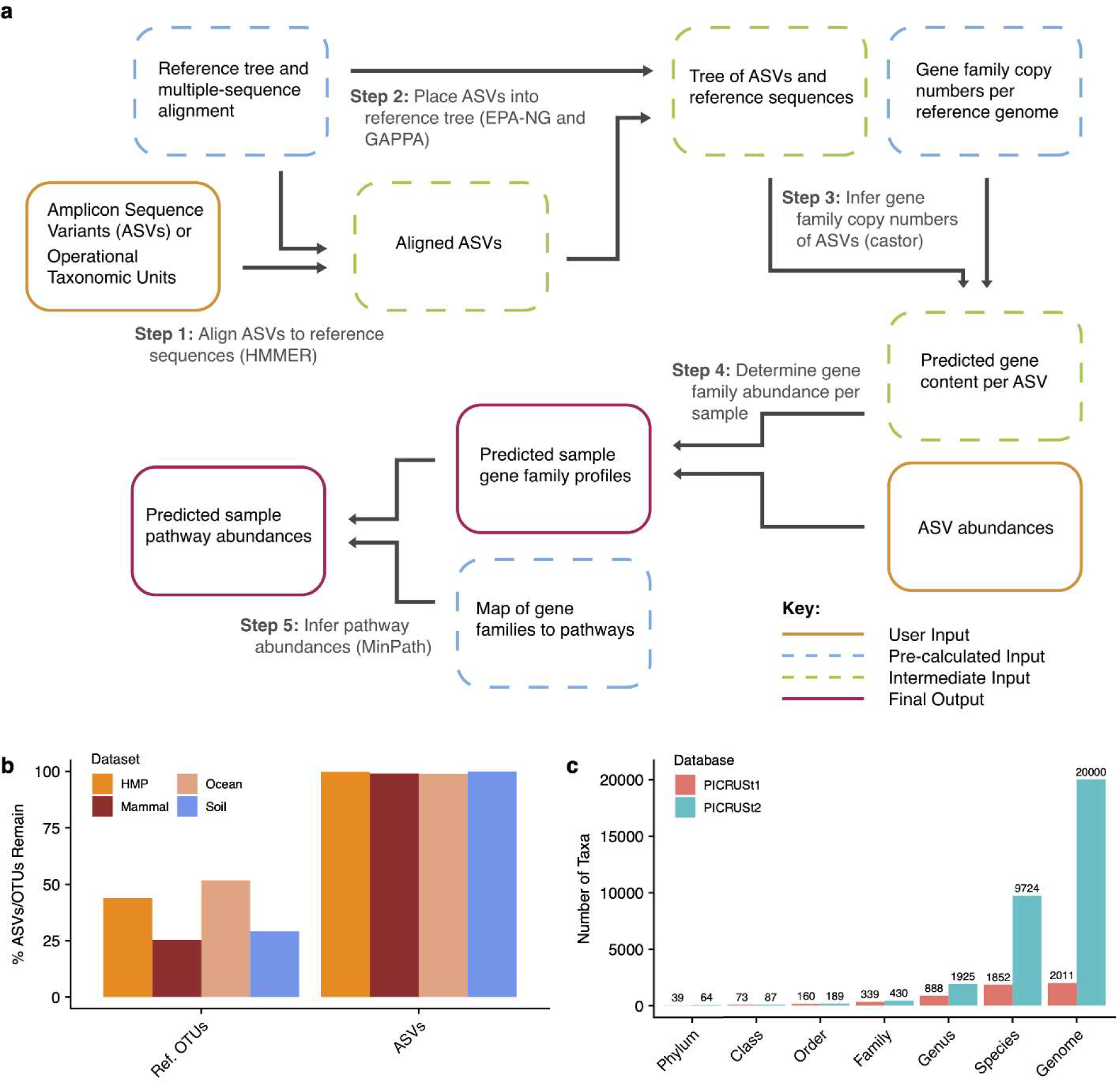
PICRUSt2 algorithm and major updates. (a) The PICRUSt2 method consists of phylogenetic placement, hidden-state-prediction, and sample-wise gene and pathway abundance tabulation. ASV sequences and abundances are taken as input, and gene family and pathway abundances are output. All necessary reference tree and trait databases for the default workflow are included in the PICRUSt2 implementation. (b) The default PICRUSt1 pipeline restricted predictions to reference operational taxonomic units (Ref. OTUs) within the Greengenes database. This requirement resulted in the exclusion of many study sequences across four representative 16S rRNA gene sequencing datasets. In contrast, PICRUSt2 relaxes this requirement and is agnostic to whether the input sequences are within a reference or not, which results in almost all of the input amplicon sequence variants (ASVs) being retained in the final output. (c) A drastic increase in the taxonomic diversity within the default PICRUSt2 database is observed compared to PICRUSt1.

PICRUSt2 integrates multiple high-throughput, open-source tools to predict the genomes of environmentally sampled 16S rRNA gene sequences. ASVs are placed into a reference tree, which is used as the basis of functional predictions. This reference tree contains 20,000 full 16S rRNA genes from prokaryotic genomes in the Integrated Microbial Genomes (IMG) database^9^. Phylogenetic placement in PICRUSt2 is based on running three tools: HMMER (www.hmmer.org) to place ASVs, EPA-ng^10^ to determine the optimal position of these placed ASVs in a reference phylogeny, and GAPPA^11^ to output a new tree incorporating the ASV placements. This results in a phylogenetic tree containing both reference genomes and environmentally sampled organisms, which is used to predict individual gene family copy numbers for each ASV. This procedure is re-run for each input dataset, allowing users to utilize custom reference databases as needed, including those that may be optimized for the study of specific microbial niches.

As in PICRUSt1, hidden state prediction (HSP) approaches are used in PICRUSt2 to infer the genomic content of sampled sequences. The castor R package^12^, which is substantially faster than the ape package^13^ used previously in PICRUSt1, now performs the core HSP functions. As in PICRUSt1, ASVs are corrected by their 16S rRNA gene copy number and then multiplied by their functional predictions to produce a predicted metagenome. PICRUSt2 also provides the ASV contribution of each predicted function allowing for taxonomy-informed statistical analyses to be conducted. Lastly, pathway abundances are now inferred based on structured pathway mappings, which are more conservative than the bag-of-genes approach previously used in PICRUSt1.

The new PICRUSt2 default genome database is based on 41,926 bacterial and archaeal genomes from the IMG database^9^ as of November 8, 2017, which is a >20-fold increase over the 2,011 IMG genomes used for PICRUSt1 predictions. Many of these genomes are from strains of the same species and have identical 16S rRNA genes. We de-replicated the identical 16S rRNA genes across these genomes, which resulted in 20,000 final 16S rRNA gene clusters.

As a result of this increased database size, the taxonomic diversity of the PICRUSt2 reference database has markedly increased compared to PICRUSt1 (**Fig. 1c**). The clearest increases in diversity have been driven by increases at the species and genus levels (5.3-fold and 2.2-fold increases respectively). However, all taxonomic levels exhibited increased diversity, including the phylum level where the coverage increased from 39 to 64 phyla (1.6-fold increase).

PICRUSt2 predictions based on the following gene family databases are supported by default: Kyoto Encyclopedia of Genes and Genomes^14^ (KEGG) orthologs (KO), Enzyme Classification numbers (EC numbers), Clusters of Orthologous Genes^15^ (COGs), Protein families^16^ (Pfam) and The Institute for Genomic Research’s database of protein FAMilies^17^ (TIGRFAM) (**Supp Table 1**). PICRUSt2 distinctly improves on PICRUSt1 by including gene families more recently added to the KEGG database. Specifically, the total number of KOs has now increased from 6,911 to 10,543 (1.5-fold increase) in PICRUSt2 compared to PICRUSt1.

We validated PICRUSt2 metagenome predictions using samples from seven published datasets that have been profiled both by 16S rRNA marker gene sequencing and shotgun metagenomics sequencing (MGS). These included three human-associated microbiome datasets: 57 stool samples from Cameroonian individuals^18,19^, 91 stool samples from Indian individuals^20^, and 137 samples spanning the human body (from the Human Microbiome Project^21^ [HMP]). These validation datasets also included non-human associated environments, including: 77 non-human primate stool samples^22^, eight mammalian stool samples^23^, six ocean samples^24^, and 22 bulk soil and blueberry rhizosphere samples^25^. These datasets span varying degrees of challenge for accurate metagenome inference due to environmental and technical factors (**Supp Table 2**).

We generated PICRUSt2 KO predictions from 16S rRNA marker gene data for each dataset. We compared these predictions to KO relative abundances profiled from the corresponding MGS metagenomes, which served as a gold-standard to evaluate prediction performance. We performed the same analysis with four alternative functional prediction pipelines: PICRUSt1, Piphillin, PanFP, and Tax4Fun2. We calculated Spearman correlation coefficients (hereafter “correlations”) for matching samples between the predicted KO abundance and MGS KO abundance tables after filtering all tables to the 6,220 KOs that could be output by all tested databases (**Fig 2**). The correlation metric represents the similarity in rank ordering of KO abundances between the predicted and observed data. The correlations based on PICRUSt2 KO predictions ranged from a mean of 0.79 (standard deviation [sd] = 0.028; primate stool) to 0.88 (sd = 0.019; Cameroonian stool dataset). For all seven datasets, PICRUSt2 predictions either performed best or were comparable to the best prediction method (paired-sample, two-tailed Wilcoxon tests [PTW] P < 0.05). Correlations based on PICRUSt2 predictions were notably higher for non-human associated datasets. This result could indicate an advantage of phylogenetic-based methods over non-phylogenetic-based methods, such as Piphillin, for environments poorly represented by reference genomes.

**Figure 2:**
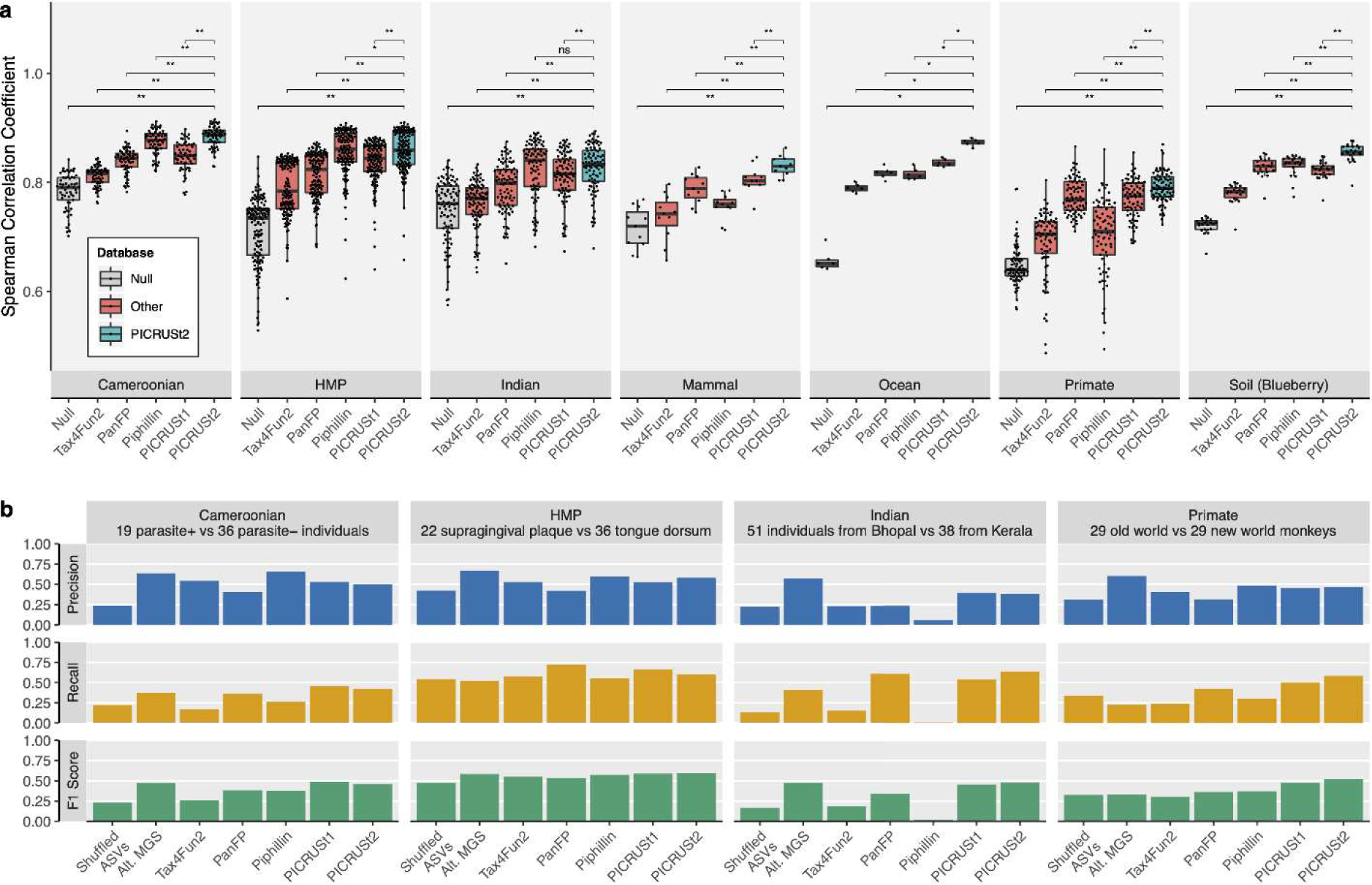
PICRUSt2 performs best or is comparable to other tools based on Spearman correlation coefficients and differential abundance results. Validation results of PICRUSt2 KEGG ortholog (KO) predictions comparing metagenome prediction performance against gold-standard shotgun metagenomic sequencing (MGS). (a) Boxplots represent medians and interquartile ranges of Spearman correlation coefficients observed in stool samples from Cameroonian individuals (n=57), the human microbiome project (HMP, n=137), stool samples from Indian individuals (n=91), non-human primate stool samples (n=77), mammalian stool (n=8), ocean water (n=6), and blueberry soil (n=22) datasets. The significance of paired-sample, two-tailed Wilcoxon tests is indicated above each tested grouping (*, **, and ns correspond to P < 0.05, P < 0.001, and not significant respectively). Note that the y-axis is truncated below 0.5 rather than 0 to better visualize small differences between categories. (b) Comparison of significantly differentially abundant KOs between predicted metagenomes and MGS. Precision, recall, and F1 score are reported for each category compared to the MGS data. Precision corresponds to the proportion of significant KOs for that category also significant in the MGS data. Recall corresponds to the proportion of significant KOs in the MGS data also significant for that category. The F1 score is the harmonic mean of these metrics. The subsets of the four datasets tested (which were the only ones with adequate sample sizes for this analysis) and the sample groupings compared are indicated above each panel. The parasite referred to for the Cameroonian dataset is *Entamoeba*. Wilcoxon tests were performed on the KO relative abundances after normalizing by the median number of universal single-copy genes in each sample. Significance was defined as a false discovery rate < 0.05. The “shuffled ASVs” category corresponds to PICRUSt2 predictions with ASV labels shuffled across a dataset (see Supplementary Text). The “Alt. MGS” category corresponds to an alternative MGS processing pipeline where reads were aligned directly to the KEGG database rather than the default HUMAnN2 pipeline.

Gene families regularly co-occur within genomes, and so the use of correlations to assess gene-table similarity may be limited by the lack of independence of gene families within a sample (**Supp Fig 2**). To address this dependency, we compared the observed correlations between paired MGS and predicted metagenomes to correlations between MGS functions and a null reference genome, comprised of the mean gene family abundance across all reference genomes. For all datasets, PICRUSt2 metagenome tables were more similar to MGS values than the null (**Fig 2a**). However, this increase over the null expectation is predominately driven by each dataset’s predicted genome content (rather than that of individual samples). This is demonstrated by the fact that these correlations are actually only slightly significantly higher than those observed when ASV labels are shuffled within a dataset (**Supp Fig 3**). The observed correlations for the shuffled ASVs ranged from a mean of 0.77 (sd = 0.196; primate stool) to 0.84 (sd = 0.178; blueberry rhizosphere). Biologically these results are consistent with several patterns. First, gene families are correlated in copy number across diverse taxa (as captured by the ‘Null’ dataset). Second, these correlations are stronger within than between environments (as shown by the difference between the ‘Null’ and ‘Shuffled ASV’ results). Lastly, environment-to-environment differences tend to be larger than sample-to-sample differences within an environment (as shown by the differences between PICRUSt2 predictions and the ‘Shuffled ASV’ results).

A complementary approach for validating metagenome predictions is to compare the results of differential abundance tests on 16S-predicted metagenomes to MGS data. A recent analysis of Piphillin suggested that this tool out-performs PICRUSt2 based on this approach^26^. We similarly performed this evaluation on the KO predictions for four validation datasets (**Fig 2b**; see Supplementary Text). Overall, PICRUSt2 displayed the highest F1 score, the harmonic mean of precision and recall, compared to other prediction methods (ranging from 0.46-0.59; mean=0.51; sd=0.06). However, all prediction tools displayed relatively low precision, the proportion of significant KOs that were also significant in the MGS data. In particular, precision ranged from 0.38-0.58 (mean=0.48; sd=0.08) for PICRUSt2 and 0.06-0.66 (mean=0.45; sd=0.27) for Piphillin. In all cases, PICRUSt2 predictions out-performed ASV-shuffled predictions, which ranged in precision from 0.22-0.42 (mean=0.30; sd=0.09). In addition, differential abundance tests performed on MGS-derived KOs from an alternative MGS-processing workflow resulted in only marginally higher precision (ranging from 0.57-0.67; mean=0.62; sd=0.04). Taken together, these results highlight the difficulty of reproducing microbial functional biomarkers with both predicted and actual metagenomics data.

MetaCyc pathway abundances are now the main high-level predictions output by PICRUSt2 by default. The MetaCyc database is an open-source alternative to KEGG and is also a major focus of the widely-used metagenomics functional profiler, HUMAnN2^27^. MetaCyc pathway abundances are calculated in PICRUSt2 through structured mappings of EC gene families to pathways. These pathway predictions performed better than the null distribution for all metrics overall (PTW P < 0.05; **Fig 3a** and **Supp Fig 4-5**) compared to MGS-derived pathways. Similar to our previous analysis, shuffled ASV predictions representing overall functional structure within each dataset accounted for the majority of this signal (**Supp Fig 4**). In addition, differential abundance tests on these pathways showed high variability in F1 scores across datasets and statistical methods with the ASV shuffled predictions contributing the majority of this signal (**Supp Fig 6**; F1 scores ranged from 0.23-0.62 (mean=0.41; sd=0.17) and 0.22-0.60 (mean=0.34; sd=0.18) for the observed and ASV shuffled PICRUSt2 predictions, respectively). Again, these results suggest that identifying robust differentially abundant metagenome-wide pathways is difficult and highlights the challenge of analyzing microbial pathways in general.

**Figure 3:**
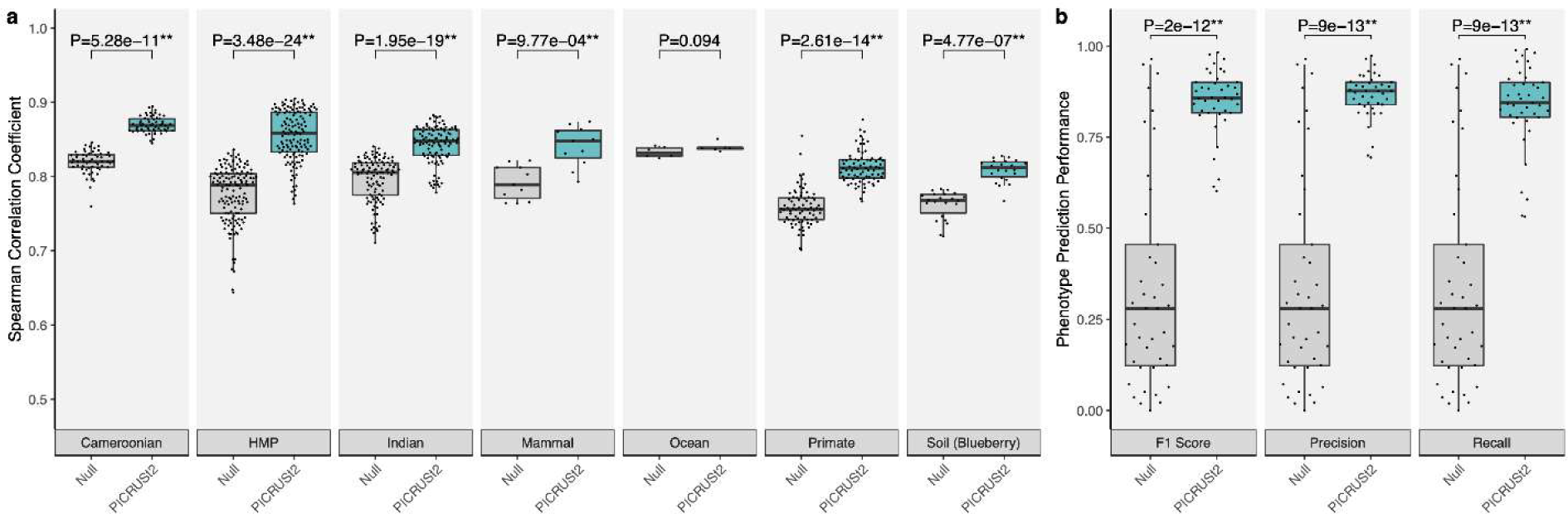
PICRUSt2 accurately predicts MetaCyc pathways and phenotypes for characterizing overall environments. (a) Spearman correlation coefficients between PICRUSt2 predicted pathway abundances and gold-standard metagenomic sequencing (MGS). Results are shown for each validation dataset: stool from Cameroonian individuals, The Human Microbiome Project (HMP), stool from Indian individuals, mammalian stool, ocean water, non-human primate stool, and blueberry soil. These results are limited to the 575 pathways that could potentially be identified by PICRUSt2 and HUMAnN2. (b) Performance of binary phenotype predictions based on three metrics: F1 score, precision, and recall. Each point corresponds to one of the 41 phenotypes tested. Predictions assessed here are based on holding out each genome individually, predicting the phenotypes for that holdout genome, and comparing the predicted and observed values. The null distribution in this case is based on randomizing the phenotypes across the reference genomes and comparing to the actual values, which results in the same output for all three metrics. The P-values of paired-sample, two-tailed Wilcoxon tests is indicated above each tested grouping (* and ** correspond to P < 0.05 and P < 0.001, respectively). Note that in panel a the y-axis is truncated below 0.5 rather than 0 to better visualize small differences between categories. The sample sizes in panel a are 57 (Cameroonian), 137 (HMP), 91 (Indian), 8 (mammal), 6 (ocean), 77 (primate), and 22 (soil).

Predictions for 41 microbial phenotypes, which are linked to IMG genomes^28^, can also now be generated with PICRUSt2. These represent high-level microbial metabolic activities such as “Glucose utilizing” and “Denitrifier” that are annotated as present or absent within each reference genome. Use of this database was motivated by the predictions made by the tools FAPROTAX^29^ and Bugbase^30^. We performed a hold-out validation to assess the performance of PICRUSt2 phenotype predictions, which involved comparing the binary phenotype predictions to the expected phenotypes for each reference genome. Based on F1 score (mean=84.8%; sd=9.01%), precision (mean=86.5%; sd=6.21%), and recall (mean=83.5%; sd=11.4%), these predictions performed significantly better than the null expectation (**Fig 3b**; Wilcoxon tests P < 0.05).

There are two major criticisms of amplicon-based functional prediction. First, the predictions are biased towards existing reference genomes, which means that rare environment-specific functions are less likely to be identified. This limitation will be partially addressed as the number of high-quality available genomes continues to grow. Moreover, PICRUSt2 allows user-specified genomes to be used for generating predictions, which provides a flexible framework for studying particular environments. The second major criticism is that amplicon-based predictions cannot provide resolution to distinguish strain-specific functionality within the same species. This is an important limitation of PICRUSt2 and any amplicon-based analysis, which can only differentiate taxa to the degree they differ at the amplified marker gene sequence.

In summary, PICRUSt2 is a more flexible and accurate method for performing marker gene metagenome inference. We have highlighted the improved performance of PICRUSt2 compared to other metagenome inference methods while also describing limitations with identifying consistent differentially abundant functions in microbiome studies. We hope that the expanded functionality of PICRUSt2 will continue to allow researchers to identify potentially novel insights into functional microbial ecology from amplicon sequencing profiles.

## Code and data availability

PICRUSt2 is available at: https://github.com/picrust/picrust2. The Python and R code used for the analyses and database construction described in this paper are available online at https://github.com/gavinmdouglas/picrust2_manuscript. This repository also includes the processed datafiles that can be used to re-generate the findings in this paper. The accessions for all sequencing data used in this study are listed in the supplementary information.

## Acknowledgements

We would like to thank Zhenjiang Xu and Amy Chen for providing us access to datafiles used for testing and the default reference database. We would also like to thank Heather McIntosh for her help designing the pipeline flowchart. GMD is funded by a Natural Sciences and Engineering Research Council (NSERC) Alexander Graham Bell Graduate Scholarship (Doctoral). VM is funded by an NIH/NIAAA Ruth L. Kirschstein National Research Service Award (F30 AA026527). MGIL is funded by an NSERC Discovery Grant and an NSERC Collaborative Research Development with co-funding from GlaxoSmithKline to MGIL and JB. CH is funded in part by NIH NIDDK grants U54DK102557 and R24DK110499. SYN is funded by an NSERC Discovery Grant.

