## Supplemental Material for "PICRUSt2: An improved and customizable approach for metagenome inference"

### Methods

#### Data availability

The raw sequencing reads analyzed in this study are available from the following online repositories. The HMP raw data is available from https://www.hmpdacc.org/HMIWGS/healthy/. The mammalian stool sequencing data is available from the Short Read Archive (SRA) under accessions SRP115632 (MGS) and SRP115643 (16S rRNA gene). The ocean sample sequencing data is available from SRA project SRP056891. The blueberry soil samples are available at SRA project accessions PRJNA484230 (MGS) and PRJNA389786 (16S rRNA gene). The Cameroonian MGS data is available under European Nucleotide Archive (ENA) project PRJEB27005 and the 16S rRNA gene sequencing data is available at MG-RAST under accession mgp15238. All of the Indian sequencing data is available under ENA project PRJNA397112. The primate metagenomics data and 16S rRNA gene sequencing data (with processed outputs) are available in the QIITA repository under accession 11212.

______The blueberry 18S rRNA gene sequencing data are available under SRA project accessions PRJNA391782 (soil) and PRJNA434067 (root). The matching MGS data for these blueberry root and soil samples are available under accession PRJNA484230. The processed ITS output files for the wine fermentation dataset are available as part of a GigaDB dataset (http://gigadb.org/dataset/100309), and the raw MGS data are available as part of SRA project accession PRJNA305659.

#### PICRUSt pipeline updates

The analyses in this paper are based on PICRUSt2 version 2.1.0-b. In addition to the improvements reported in the main text, several other updates have also been made to the

PICRUSt pipeline. Since the HSP step is now run using the castor R package, other inference approaches like maximum parsimony (MP) may be performed in realistic time-frames besides phylogenetic independent contrasts[1], which was the default approach in PICRUSt1. The default HSP method is now MP with a parameter weighting the contribution of branch lengths set to 0.5 (*edge_exponent* option in castor package). This parameter value was chosen because setting this parameter to a non-zero value resulted in more reproducible predictions.

In addition, now that any study sequences can be input to PICRUSt, and not just Greengenes closed-reference OTUs, a nearest-sequence taxon index (NSTI) screening step is recommended to eliminate sequences above a certain cut-off. The default NSTI cut-off in PICRUSt2 is two, which was chosen as an extremely lenient cut-off intended to eliminate problematic sequences. Only one ASV in the test inflammatory bowel disease dataset (see below) was above this cut-off, which corresponded to a mitochondrial sequence. The only sequences above this cut-off in the HMP validation dataset corresponded to two 18S ASVs that were clustered within the 16S rRNA gene dataset. Similarly, although 13/1148 of ASVs in the ocean dataset were above the NSTI cut-off of two, these ASVs corresponded to candidate taxonomic groups that have no representative reference genomes in the default PICRUSt2 database. Based on these observations, we believe this cut-off should be suitable for most scenarios; however, users can select a NSTI value that best fits their study design and environment (i.e. whether to maximize precision or recall).

Transforming gene family predictions to pathway abundances in PICRUSt1 was done by assuming that the abundance of each gene family contributed equal abundance to all pathways containing the gene family (i.e. if a gene family can be involved in 10 pathways the gene family abundance would be added equally to the abundance of all 10 pathways). Although this approach is easy to understand, it results in a high false-positive rate of identifying pathways present. To improve on this approach, we adapted the approach taken by HUMAnN2[2] v0.11.1 into the PICRUSt2 pipeline. MinPath[3] (v1.2 as modified for the HMP workflow[4]) is first run to identify the minimum pathways present given the gene families present. By default, these predictions are made based on the EC number predictions after regrouping them to MetaCyc reactions to predict MetaCyc pathway abundances. The mappings files and code for regrouping to MetaCyc reactions and mapping from reactions to structured pathways were taken from HUMAnN2. We further split the pathway mapping files into prokaryotic and fungal sets based on the taxa where these pathways have been identified (as reported in the MetaCyc online database).

16S rRNA gene database processing

The 16S rRNA gene sequences and gene family counts from a total of 52,217 genomes were acquired from IMG on 8 Nov. 2017. These data were based on IMG annotations and we did not work with the raw genome sequences. Genomes lacking a 16S rRNA gene length of at least 1,250 bp or that were identified as eukaryotic marker genes were removed. The gene families in these annotations corresponded to these databases: KEGG (v77.1), Pfam (v30), TIGRFAM (v15), COG (v2014), EC numbers (as of 21 Jan 2016).

We also created an alternative trait database containing phenotypes defined by IMG[5]. These phenotypes are more directly interpretable than the gene family databases described above. Files listing IMG genome ids positives for one of 65 phenotypes were downloaded on 8 Jan 2019. The presence and absence of phenotypes was re-coded as 1 and 0. Prototrophic and auxotrophic phenotypes for the same compound were combined into a single prototrophic phenotype (auxotrophs are coded as 0 and unknown phenotypes are coded as NA). After this merging step, and removing two extremely rare phenotypes, there was a final set of 41 phenotypes remaining.

To identify low-quality, incomplete genomes, and possible misassembled assemblies we calculated the median number of single-copy KEGG orthologs (KO) as previously identified for the tool MUSiCC[6]. Since these genes are expected to be found in single copies within each genome, we reasoned that incomplete or contaminated genomes could be identified by a median copy number less or greater than one, respectively. Accordingly, we discarded all genomes with a median number of single-copy genes that differed from one. In addition, we discarded genomes lacking a sufficient number of genes within any gene family. The minimum number of gene families per genome was chosen based on visualizing the distribution of gene family numbers over all genomes and choosing a cut-off that eliminated outliers (minimum cut-offs of unique gene families were 500, 250, 500, 750, and 350 for the COG, EC, KO, Pfam, and TIGRFAM databases). Importantly, this filtering means that endosymbionts and other organisms with reduced genome sizes will be underrepresented in the PICRUSt2 reference database. After these filtering steps, a total of 10,291 genomes were discarded, producing a final set of 41,926 genomes. Gene family copy numbers higher than 10 in this final set were re-coded to be 10 to decrease the number of possible prediction states.

Since prokaryotes often have multiple 16S rRNA gene copies, the centroid 16S rRNA gene per genome was identified using the VSEARCH[7] (v2.4.4) *cluster-fast* command with an identity cut-off of 90%. In cases where multiple centroid sequences were found, a single centroid was chosen randomly. There were 3,002 such clusters and the sequences contained in these clusters made up 59.5% of the original 41,926 genomes. Among the 20,000 sequence clusters, there was a mean of 2.1 sequences per cluster (standard deviation = 562.5). The cluster with the highest sequence count of 1,379 corresponded to strains of *Staphylococcus aureus*. We observed a mean clustered-sequence length of 1489.6 base-pairs (bp; standard deviation = 65.8) overall.

The final 16S rRNA gene sequences were used to build a multiple sequence alignment (MSA) using ssu-align (v0.1.1; http://eddylab.org/software/ssu-align/) against the bacteria alignment model. Note that although the majority of the reference 16S rRNA gene sequences correspond to bacteria, there are archaeal sequences as well, which the bacteria ssu-align alignment model is likely less appropriate. Only weak masking of this output MSA was performed (*ssu-mask* options: *--pf 0.001 --pt 0*). The custom Python script *derep_fasta.py* was used to identify sequences in this alignment after this masking step. A phylogenetic tree was built from this MSA with RAxML-ng[8] (v0.6.0) using the *GTR+G* model. The custom Python script *mean_16S_function_counts.py* was used to calculate the mean gene family abundances for all sequences within a cluster. These values were then rounded to the nearest integer. The full NCBI taxonomic lineage of all 16S rRNA gene clusters was called by the taxizedb R package (https://github.com/ropensci/taxizedb) using the species name provided by the IMG FASTA metadata. Gene family trait depths were calculated using the castor R package function *get_trait_depth* with default settings, which is based on the consenTRAIT metric[9].

#### Amplicon dataset processing

An in-depth comparison of the technologies and sequencing depths for each of the seven 16S rRNA gene validation datasets is shown in Supplementary Table 2. The processing pipelines and filtering criteria differed for each dataset due to technical differences between them. The key difference was that DADA2[10] was run for the HMP dataset because this was Roche 454 sequence data. Deblur[11] was run on all other validation datasets since they were Illumina sequence data (Supp Table 2).

The HMP reads were filtered using DADA2 (v1.6.0) options. Denoising these sequences with DADA2 resulted in 1,865 ASVs after discarding ASVs with a minimum frequency of 10 and discarding 17 samples with fewer than 2,000 reads. The reverse complement of these sequences was taken before running the functional prediction pipelines. The mammalian stool dataset was run using deblur with the default options in QIIME 2[12] (v2017.12), which resulted in 323 ASVs after discarding two samples with final read counts less than 3,220. The Cameroonian, Indian, ocean, and blueberry soil datasets were also processed with deblur and no post-processing was done (besides discarding samples that did not overlap with the MGS data) because all samples had high depth, which resulted in 4,077, 2,237, 1,148, and 3,333 ASVs for each dataset respectively. The primate dataset previously processed by deblur was acquired through the QIITA database[13], which contained 7,452 ASVs after excluding samples with no MGS data available.

Before running PICRUSt1 on each dataset, ASVs were matched against the Greengenes v13_8 OTUs using the VSEARCH command *–usearch_global* with an identity cut-off of 97%. The ASVs were then regrouped to be the best matching Greengenes OTU to be compatible with the default PICRUSt1 pipeline.

The alternative functional prediction tools discussed in this paper were the following versions (with default reference databases): PICRUSt version 1.1.3, Tax4Fun2 version 1.1.3, PanFP from GitHub commit: 1f49bd1b7341b47d46fa7eaa45d7771044d0efde, Piphillin online interface as of 7th Nov. 2019 at http://piphillin.secondgenome.com (KEGG v88.1) and PAPRICA version 0.5.2.

We also created several shuffled prediction datasets to help evaluate the PICRUSt2 predictions. These shuffled datasets were based on shuffling the predicted genome content and not the relative abundance of the ASVs across samples. This shuffling was performed on entire predicted genomes, i.e. each ASV was assigned the genomic content of a randomly sampled ASV (see below). This approach allowed us to assess how randomizing predicted genomic content for ASVs across samples in a dataset affects the concordance with MGS data. These shuffled datasets also provide a baseline of the performance expected by chance for the differential abundance validations. In other words, they give a baseline of the expected concordance (e.g. precision and recall) based on differential abundance testing compared to MGS data given the same ASV relative abundance across samples but shuffled predicted genomes.

More specifically, these datasets were produced by shuffling the ASV ids in each PICRUSt2 prediction table (i.e. the first column of the PICRUSt2 output prediction tables). This was performed 10 times for each dataset and then averaged to produce a single shuffled table per dataset. Shuffled MetaCyc pathway abundances were produced by running the PICRUSt2 pathway pipeline on the shuffled EC metagenome tables. These shuffled datasets are referred to as the "Shuffled ASVs" category in the main text and figures.

#### Shotgun metagenomics sequencing validation dataset processing

All shotgun metagenomic sequencing (MGS) datasets were processed using the same pipeline, which is described below. Each dataset was filtered using kneaddata (v0.6.1; https://bitbucket.org/biobakery/kneaddata/wiki/Home) to run (1) Trimmomatic[14] (v0.36) to exclude low-quality reads with the options *SLIDINGWINDOW:4:20* and *MINLEN:50* and (2) bowtie2[15] (v2.3.2) to exclude reads that mapped to the human and PhiX genomes with the options *–very-sensitive* and *--dovetail*. For the blueberry-associated samples we also mapped reads against the northern highbush blueberry (*Vaccinium corymbosum)* genome[16] version W8520 (downloaded from https://www.vaccinium.org on October 29, 2018) to exclude additional contaminant reads. For samples with paired-end reads the forward and reverse reads were concatenated into the same file. HUMAnN2 was then run to identify the abundances of annotated UniRef50 gene families in each sample. The abundances of gene families were regrouped into other gene family databases as indicated in the text using *humann2_regroup_table*. Pathway abundances and coverages were produced by the default HUMAnN2 mapping files. These steps were parallelized when possible with GNU Parallel[17] (v20170722).

To compare how different metagenomics processing pipelines can affect the resulting functional abundance tables, we also ran HUMAnN2 directly against the KEGG v56 database using default options. This pipeline only involved mapping translated reads against this database (i.e. it skipped the nucleotide alignment step) and results in KO abundances directly for each sample. This differs from the other pipeline described above, because in the above case UniRef50 ids were regrouped to KOs based on a mapping file rather than based on direct read mappings. These KO abundances are referred to as the alternative MGS pipeline ("Alt. MGS") in the main text and figures.

16S rRNA gene-MGS validation analyses

The simulation approach to illustrate the issue of high null Spearman correlation coefficients was based on the following procedure. First, for each *N* in the set of integers that span 1 to 100, two subsets of *N* genomes were sampled randomly from the reference database. The abundance of each sampled genome was taken from a negative binomial distribution family implemented in the R function, *rnbinom*, with parameters *size=10* and *prob=0.7*, which aimed to simulate the over-dispersed count data frequently encountered in sequenced data sets. Gene family abundance tables were then computed for each of the two subsets of genomes based on the abundance of each genome and the abundances of gene families within each genome. Spearman correlation coefficients were then computed between the two gene family tables. This procedure was replicated 10 times for each *N*.

This simulation inspired the use of null distributions in our validation analyses. We calculated the correlations between the MGS metagenome or gene table and a synthetic gene table comprised of the mean gene count number across all reference genomes in the database, which is referred to as the "null expectation" through-out the main-text. The null Spearman correlation coefficient distributions of pathway abundances and coverages were similarly based on the reference genome pathways inferred from the EC reference database.

For the purposes of comparing functional prediction tools, gene family tables were filtered to only those gene families present in the databases of all tested functional prediction tools. Gene families absent in all samples were retained as zeros and were not removed. We converted the predictions to binary presence (positive) and absence (negative) format before calculating precision and recall (any abundance greater than zero was considered as evidence for presence). Precision and recall are defined as $TP/\,(TP\, +\, FP)$ and $TP/\,(TP +\, FN)$, respectively, where TP=number of true positives (i.e. functions correctly called as present), FP=number of false positives, TN=number of true negatives, and FN=number of false negatives. The F1 score is the harmonic mean of precision and recall: 2 * ((precision * recall) / (precision + recall)).

We also conducted differential abundance tests to compare how results differ between predicted metagenomes and actual MGS data. To conduct these analyses, we subset four of the validation datasets into sample groupings appropriate for pairwise testing. The other datasets were excluded because they resulted in no significant results based on the MGS data after restricting the data to appropriate sample groupings. The comparisons were:

- Stool samples from 19 *Entamoeba*-positive vs 36 *Entamoeba*-negative Cameroonian individuals
- 22 supragingival plaque vs 36 tongue dorsum samples from the Human Microbiome Project (i.e. samples from two different oral body sites).
- Stool samples of 51 individuals from Bhopal, India vs 38 individuals from Kerala, India
- Stool samples of 29 old world monkeys vs 29 new world monkeys

In the main text the differential abundance results are focused on Wilcoxon tests on relative abundance values for the KOs (after normalizing function abundances by the median number of universal single-copy genes per sample[6]). The pathway differential abundance results are also based on Wilcoxon tests, but on the relative abundance of pathways instead.

As a comparison point we also ran ALDEx2[18] (Wilcoxon test with 128 Monte Carlo samples) and DESeq2[19] to see how these choices affect the resulting significant functions. DESeq2 was run twice on each dataset: once with default options and separately with options specifically recommended for microbiome data. In the latter case, these options included first calculating the geometric means of the data to estimate size factors and then using the option fitType="local". This second method is referred to as "DESeq2 GMeans" in Additional File 1.

We also tested for differential prevalence of functions based on the presence and absence of functions with Fisher's exact tests.

ALDEx2 is a general method for compositional data analysis that used a Dirichlet-multinomial model to infer sampling and biological variation. The significance reported by this tool is based on Wilcoxon tests after abundances are estimated from the count data. DESeq2 is also a compositional data analysis method for read counts, based on the negative binomial distribution. Lastly, the Fisher's exact tests tested for differential prevalence rather than abundance. In this case, the tests were focused on the counts of how many samples were positive (i.e. had the specific function) compared to those that were negative in each group based on the binary presence/absence of functions.

ALDEx2 and DESeq2 require count tables as input, which required the prediction tables to be rounded. Tax4Fun2 and PanFP were excluded from these analyses, because they do not output predictions in a format that corresponds to the original ASV relative abundances (i.e. the output tables cannot be meaningfully rounded). In all cases significant functions were identified based on Benjamini-Hochberg corrected P-values < 0.05.

Additional notes on statistical analyses

No tests for statistical power were conducted to determine sample sizes for this study. In the boxplots throughout this paper the centre line corresponds to the median and the lower and upper hinges (i.e. the edges of the "box") represent the $25^{th}$ and $75^{th}$ percentiles, respectively. The boxplot whiskers extend to the most extreme values no further than 1.5 multiplied by the inter-quartile range in each direction. The points overlaid on the boxplots correspond to each individual sample unless otherwise stated.

#### Supplementary Methods

Inflammatory bowel disease data processing and analyses

We ran PICRUSt2 on a dataset of ileal biopsies from an inflammatory bowel disease (IBD) cohort to highlight how metagenome inferences can be generated for datasets where MGS is infeasible. Raw 16S rRNA gene reads and processed human host transcriptome and metabolome tables were downloaded from https://ibdmdb.org. The 16S rRNA gene data was processed using deblur and QIIME 2 as described above for the validation datasets. Only ASVs found in at least two samples and called by at least 10 reads were retained, which resulted in 1,419 final ASVs. The MGS raw reads were processed using the same workflow as the validation datasets described above. PICRUSt2 was run with default options except for the option --*per_sequence_contrib*, which was set to get pathway abundances within each predicted genome for ASV-specific analyses. The unstratified pathway abundances were then calculated by summing over the pathways contributed by each ASV within each sample. Only features present in at least 33% of samples were retained for all analyses. In addition, any pathways described as "superpathways" or "engineered" were excluded from these analyses. ALDEx2[18] (v1.12.0) was run with default options to identify features at differential relative abundance between Crohn's disease and control subjects for taxa and pathways independently.

Partial Spearman correlations between predicted pathway abundances and both the metabolomic and transcriptome data was conducted with the R package ppcor (v1.1). Subject consent age was controlled for when calculating the partial correlations. Before calculating these correlations, pathway abundance data was first transformed by the arcsine square-root transformation, and the metabolomic and transcriptomic datasets were transformed by $\log_{10}$ after adding a pseudocount of 1. Metabolites were limited to those with non-empty compound names and the gene expression data was limited to 11 genes known to be biomarkers of CD-associated ileal inflammation[20]: DUOXA2, MMP3, AQP9, IL8, DUOX2, APOA1, NAT8, AGXT2, CUBN, FAM151A, and NOD2. Because several of these genes are highly correlated, we removed redundant genes based on hierarchical clustering of the complement of Spearman correlation coefficients between all genes. Six clear clusters of genes were then identified and the following six representative genes for each cluster were retained for further analyses (the other genes in each cluster are indicated in parentheses): DUOX2 (DUOXA2), MMP3, AQP9 (IL8), APOA1, NAT8 (AGXT2, CUBN, FAM151A), and NOD2.

18S rRNA gene and ITS database processing

A total of 574 publicly available fungi genomes were downloaded from the 1000 Fungal Genomes Database (http://1000.fungalgenomes.org) on November 16, 2018. The 18S rRNA genes were annotated using barrnap (v0.9-dev; https://github.com/tseemann/barrnap), and 18S rRNA genes were parsed from the genomes using the custom Python script *rRNA_from_gff3.py*. ITS sequences were identified and parsed from all genomes using ITSx[21] (v1.0.11) using the --*only_full T* and *–heuristics* options. Sequence length cut-offs for the 18S rRNA genes and ITS sequences were 605-3,076 bp and 146-2,570 bp, respectively. BUSCO[22] (3.0.2) was run to identify incomplete and contaminated genomes with the *fungi_odb9* database. Only the genomes with a completeness of at least 70%, and a duplicated metric (which is based on the copy number of single-copy genes) of at maximum 10% were retained. After restricting genomes to those that passed these quality cut-offs that also had at least one passing amplicon region, there were 229 genomes in the 18S rRNA gene database and 201 genomes in the ITS database.

The 18S rRNA gene and ITS sequences were then dereplicated using the same approach as used for the 16S rRNA genes. The 18S rRNA gene MSA was built using the ssu-align pipeline as for the prokaryotic database (using the *eukarya* alignment model) whereas the ITS MSA was built using MAFFT[23] (v7.407) with the *–genafpair* and *–maxiterate 1000* options. Phylogenetic trees for both MSAs were built with RAxML-ng (v0.8.0) as for the prokaryotic database except a guide tree enforcing a taxonomic topology was also used. EC number copy numbers per genome were downloaded for these genomes also from the 1000 Fungal Genomes Database. Mean EC number abundances were calculated for dereplicated amplicon sequences using the same the approach implemented for the prokaryotic databases.

18S rRNA gene and ITS amplicon data processing

The blueberry soil 18S rRNA gene data was processed the same as the blueberry soil 16S rRNA gene data except the output ASVs were restricted to those classified as fungi. This resulted in a total of 1,048 ASVs and a minimum sample depth of 1,981 reads. A total of 3,691 ASVs and a minimum depth of 2,091 ASVs was produced when re-running this pipeline with all blueberry-associated 18S rRNA gene samples (i.e. including blueberry root and soil samples with no matching MGS data). The R package rfPermute (v2.1.6) was run with 501 trees, 1,000 replicates, and the default *mtry* setting to identify significantly different predicted pathways between the three environments based on PICRUSt2 predictions run on this full blueberry-associated dataset. Previously clustered ITS sequences and a processed abundance table were acquired for the wine fermentation dataset[24]. These files were used because no raw ITS reads could be located for this dataset.

When comparing these amplicon datasets with the corresponding shotgun metagenomics data, the percent of eukaryotic DNA in the MGS data was identified with Metaxa2[25] (v2.2), which parses rRNA genes from the raw reads.

#### Supplementary Results

Paired 16S rRNA gene and shotgun metagenomics validations

We previously developed the nearest sequenced taxon index (NSTI) as a metric for summarizing a microbial taxonomic profile's novelty relative to isolate genomes[26]. This measures the abundance-weighted distances in the phylogenetic tree between taxa (ASVs) from a community and the tips of the nearest sequenced neighbours. These NSTI distributions are calculated automatically in PICRUSt2 and differed significantly among the evaluation datasets, demonstrating their range of "unusualness" relative to sequenced isolates (Kruskal-Wallis $\chi 2$=19,499, $P < 2.2 \times 10^{-16}$; Supp Fig 7a and b). Datasets from more well-characterized communities have lower mean NSTI values overall as expected, ranging from 0.10 (sd: 0.11) in the Indian dataset to 0.51 (sd: 2.06) in the ocean dataset. A maximum NSTI cut-off of two is implemented by default in PICRUSt2, as a guideline to prevent unconsidered interpretation of overly speculative inferences, which resulted in a mean of 0.27% (sd: 0.4) of ASVs being excluded across these datasets. These excluded ASVs mainly correspond to either eukaryotic sequences or microbial phyla with no reference genomes available.

In addition to the Spearman correlations reported in the main text, we also investigated metagenome predictions based on the presence and absence of output KOs by calling any function with non-zero abundance as present and assessing relative differences in precision, recall, and F1 score (see methods for definitions). PICRUSt2 and Piphillin exhibited the best (or non-statistically different) F1 score across 3/7 (HMP, mammalian stool, and ocean) and 4/7 (Cameroonian stool, HMP, Indian stool, and soil datasets) datasets, respectively (Supp Fig 8); and were substantially better than the null (<0.6 in all datasets). Investigating precision and recall revealed that Piphillin tended to have higher precision scores (fewer false positives) while PICRUSt2 had better recall (fewer false negatives) (Supp Fig 9). The slightly lower precision of PICRUSt2 is driven by falsely called KOs that are on average at significantly lower relative abundance in the PICRUSt2 results compared to the Piphillin results (PTW $P < 0.05$; mean of 0.0020% [sd: 0.0015] for PICRUSt2 vs mean of 0.0055% [sd: 0.0047] for Piphillin). However, it remains unclear what proportion of these disagreements between PICRUSt2 KO predictions and the MGS data are in fact false negatives in the MGS data due to low sequencing depth or annotation limitations from short MGS reads. Indeed, there is a significant positive relationship between the number of annotated MGS reads per sample and the observed precision of PICRUSt2 predictions (Spearman correlation coefficient across all datasets combined: 0.80; $P < 2.2 \times 10^{-16}$; mean coefficient of 0.49 [sd: 0.15] for individual datasets).

We also evaluated the performance of the PICRUSt2 EC predictions by comparing with PAPRICA, which is another EC-based prediction tool. Correlations between EC predictions with the EC gene families observed in MGS data were significantly higher than PAPRICA for 4/7 validation datasets (PTW $P < 0.05$; Supp Fig 10). In addition, the recall of PICRUSt2 predictions was significantly higher than PAPRICA although this came at a cost of precision (PTW $P < 0.05$; Supp Fig 11).

The predicted MetaCyc pathways generated by PAPRICA drastically differ from the metagenomics pathway abundances (Supp Fig 12) likely because PAPRICA implements a different approach for inferring pathway abundances. For this reason, evaluating the performance of the PAPRICA pathway predictions against HUMAnN2 pathways may not be appropriate.

In addition to the differential abundance validations reported in the main text we also tested several other statistical methods. These methods included ALDEx2 and DESeq2 (with default options and options specifically recommended for microbiome data) as well as Fisher's exact tests to test for differences in prevalence between samples. The raw results of these analyses are reported in Additional File 1. The DESeq2 results with both option selections were extremely similar, so we excluded the results based on non-default DESeq2 options from subsequent analyses. Overall, the different statistical methods result in different sets of significant functions for both the MGS data and the predicted metagenomes. In addition, the F1 scores based on the different differential abundance tools can vary substantially (Supp Fig 13). Fisher's exact tests for differential prevalence between sample groupings have especially low F1 scores (ranging from 0.02-0.29 for PICRUSt2). We believe that a more in-depth comparison of these statistical tests is beyond the scope of this manuscript, but this result does nonetheless highlight the challenge of reliably reproducing biomarkers from predicted metagenomes.

This challenge is especially apparent for the MetaCyc pathways (Supp Fig 6), for which the PICRUSt2 predictions agreed only slightly better with the MGS pathways compared to the ASV shuffled datasets. The PICRUSt2 predicted pathways varied in precision and F1 score from 0.23-0.63 (mean=0.39; sd=0.19) and 0.23-0.62 (mean=0.41; sd=17), respectively. The shuffled ASV-based pathway predictions had similar performance, ranging from 0.16-0.58 (mean=0.31; sd=0.19) and 0.22-0.60 (mean=0.34; sd=0.18) for the precision and F1 scores, respectively. These values are based on running Wilcoxon tests on the relative abundances of MetaCyc pathways. The alternative differential abundance (and prevalence) statistical methods resulted in similar concordances between the observed and ASV shuffled pathway predictions (Supp Fig 6).

Functional profiling of inflammatory bowel disease using PICRUSt2

To demonstrate the utility of PICRUSt2 in making functional inferences in a human health context where only amplicon sequencing is feasible, we profiled 27 ileal biopsy samples from subjects with Crohn's disease (CD) and 20 control subjects (non-IBD). These data are a subset of the Inflammatory Bowel Disease Multi'Omics Database[27], which provides "multi-omics" data to identify host and microbial features associated with inflammatory bowel disease (IBD). Our analysis was based on 16S rRNA gene libraries collected from biopsy samples in this dataset. Importantly, MGS could not practically be performed on these (or any typical) biopsy samples due to the overwhelming predominance of human host DNA, which competes with microbial DNA for sequencing reads. As such, MGS data was produced only for subject stool samples for this study, which are analyzed here in addition to accompanying human RNA-seq transcriptional profiles (from the same biopsies) and metabolomic profiles from paired stool.

We first analyzed these data with a typical testing framework for identifying significant microbial features: testing for significantly differentially abundant ASVs clustered by taxonomy as well as inferred pathway abundances. Based on this standard approach we identified no pathways that significantly differed between CD and non-IBD subjects (FDR < 0.1). However, five taxa were identified with a differential relative abundance between non-IBD and CD subjects based on a lenient FDR q-value cut-off of 0.2. These included four taxa within the Clostridiales order, which were increased in relative abundance in control subjects, and the phylum Proteobacteria at higher relative abundance in CD subjects.

We next focused on the predicted MetaCyc pathways inferred by PICRUSt2 for ASVs underlying the significantly differentially abundant taxa: (1) 35 significant Clostridiales ASVs and (2) 192 Proteobacteria ASVs. For each predicted pathway in the community, we calculated the ratio of the abundance of that pathway contributed (i.e. potentially produced) by ASVs within the group of interest compared to the pathway's abundance contributed by all other ASVs. Based on this approach we identified three pathways significantly contributed by Proteobacteria (Supp Fig 14a; Wilcoxon test FDR $< 0.05$). In addition, the relative contribution to 78 pathways by Clostridiales significantly differed between CD and non-IBD subjects (Wilcoxon test FDR $<$ 0.05). These results demonstrate how PICRUSt2 stratified outputs allow integration of functional predictions with taxonomic findings as opposed to treating the two independently as in non-contributor stratified metagenomic analyses.

We next investigated whether analysis of stool MGS data rather than ileal PICRUSt2 predictions resulted in substantially different conclusions, either due to methodology or body site. The number of classified genera contributing to each pathway within CD subjects differed strikingly depending on whether the contributors were identified through ileal 16S rRNA gene sequencing or stool MGS (mean difference: 7.3; sd: 10.2; Supp Figure 14b). While a small number of pathways were uniquely identified by stool MGS, for most pathways a greater number of contributing taxa were identified by PICRUSt2. This is most likely due to the much greater taxonomic diversity accessible through amplicon databases than through reference genome isolates. This result could also be due to biological differences between the stool and ileal samples. However, we also identified a similar trend in the paired 16S rRNA gene-MGS datasets, although the magnitude of difference depended greatly on sequencing technology and sampling environment (Supp Figure 15). Not just the number of taxa, but also their identities, differed between stool metagenomes versus biopsy inferences. For instance, for the tetrapyrrole biosynthesis I (from glutamate) pathway (PWY-5188), the top contributors differ between phylum Proteobacteria in biopsy 16S rRNA gene profiles, while *Akkermansia* (in phylum Verrucomicrobia) is the top contributor identified in the MGS data (Supp Figure 14c).

Last, we tested whether the PICRUSt2 predictions give novel insights into CD biomarkers by associating 207 predicted pathways with both 583 metabolites from paired stool metabolomic profiles and the ileal transcription levels for six human host genes of interest. We identified no significant associations between predicted pathway and stool metabolite levels, but 29 associations between the predicted pathway and ileal transcript levels (FDR < 0.1; Supp Table 3). Some of these significant associations are driven largely by individual taxa. For example, since the predicted relative abundance of chondroitin sulfate degradation is entirely contributed by *Bacteroides* (Supp Figure 14d), the association between this pathway and NAT8 expression (partial R=-0.58) is trivially due to the relative abundance of this genus. However, not all significant associations are driven by individual taxa. For instance, there is no single taxon driving the association of the predicted relative abundance of the methylphosphonate degradation I pathway with MMP3 expression (partial R=-0.62; Supp Figure 14e). This association is an example of PICRUSt2 predictions yielding potentially novel insights beyond those of the originating amplicon-based taxonomic profiles.

<u>Validation of fungal metagenome inference</u>

We next assessed PICRUSt2's new capabilities for predicting metagenomes based on fungal amplicon sequencing of either the 18S rRNA gene or internal transcribed spacer (ITS) regions. Data used for these predictions included EC abundances from 294 fungal genomes from the 1000

Fungal Genomes Project that were publicly available as of November 16, 2018 and passed quality control criteria. Unlike the prokaryotic database, only a minority of 18S rRNA gene and ITS sequences were redundant across genomes (7.5% and 8.5%, respectively). A total of 7 and 8 phyla as well as 183 and 209 genomes are represented in the ITS and 18S rRNA gene databases, respectively (see Supp Table 4 for the database counts at all taxonomic levels).

We first evaluated the performance of PICRUSt2 18S rRNA gene and ITS metagenome predictions by leave-one-out cross-validation of individual genomes. Spearman correlations were calculated between the predicted EC abundance profiles in each held-out genome and the EC abundances in the known genome. For both the 18S rRNA gene (Spearman Rho mean=0.821; sd=0.141) and ITS databases (Spearman Rho mean=0.822; sd=0.135), the predictions were significantly better than the null expectation (Wilcoxon test $P < 0.001$; Supp Fig 16). Similar to the 16S rRNA gene-based validations, genome prediction accuracy decreased as reference genomes were artificially held out of the training dataset at increasing taxonomic scale, suggesting that overall accuracy is hampered for those lineages without comprehensive representative genomes.

Next, we evaluated the performance of fungal EC predictions on two amplicon sequencing datasets with paired MGS data using the same approach as with 16S rRNA gene predictions. The two validation datasets used were the same 22 blueberry soil samples described in the main-text, which also underwent 18S rRNA gene sequencing[28], and eight wine fermentation internal transcribed spacer (ITS1) sequencing samples[24]. EC predictions for both of these datasets were significantly more similar to the MGS gold-standard compared to the null expectation based on correlations (Supp Fig 17a; $P=9.5 \times 10^{-7}$ and $P=7.8 \times 10^{-3}$ for the blueberry soil and wine fermentation datasets respectively). However, the correlations observed for these datasets was substantially lower than for the 16S rRNA gene-based validations, which is to be expected as these metagenomes include a substantial amount of functions from non-fungal origins. The mean correlations in the blueberry soil dataset were 0.340 (sd: 0.016) and 0.365 (sd: 0.019) for the null expectation and PICRUSt2 predictions, respectively. There was a larger mean difference for the wine fermentation dataset where the mean correlations were 0.492 (sd: 0.004) and 0.611 (sd: 0.028) for the null expectation and PICRUSt2 predictions, respectively. Interestingly, the correlations based on predicted MetaCyc pathway abundances were slightly lower than the null values for the blueberry soil 0.461 (sd: 0.009) and wine fermentation 0.501 (sd: 0.008) datasets (Supp Fig 17b).

One potential factor affecting these results is the percent of eukaryotic DNA within the MGS data. A low percent of eukaryotic DNA would result in prokaryotes mainly contributing to gene family abundances. The percent of eukaryotic DNA (after excluding plant and animal DNA) within the MGS datasets differed dramatically between the blueberry soil (mean: 8.17%; sd: 5.82) and wine fermentation datasets (mean: 96.72%; sd: 1.44; Supp Fig 17c). This low percent of eukaryotic DNA in the blueberry soil dataset could partially account for the relatively poor performance we observed.

To investigate whether the PICRUSt2 predictions for the blueberry soil dataset can nonetheless distinguish sample groupings, we ran PICRUSt2 on 18S rRNA gene sequencing data from additional blueberry soil samples with no matching MGS data as well as blueberry root samples from the same sampling location[29]. We then generated a Random Forest model to identify the most informative predicted EC numbers that distinguish samples by whether they were taken from a bulk soil, rhizosphere, or root environment. This model resulted in a classification accuracy of 68%, which was substantially better than the random expectation (accuracy 36%) and identified 32 significantly informative EC numbers ($P < 0.001$; Supp Fig 17d). Importantly, although this model was more accurate than random, this analysis does not prove that the predicted EC numbers themselves were accurately predicted. Instead, these combined analyses demonstrate a proof-of-concept that fungal metagenomes can be predicted more accurately than expected by chance. However, due to the low correlation values we observed, these predictions are unlikely to provide reliable biological insights in practice.

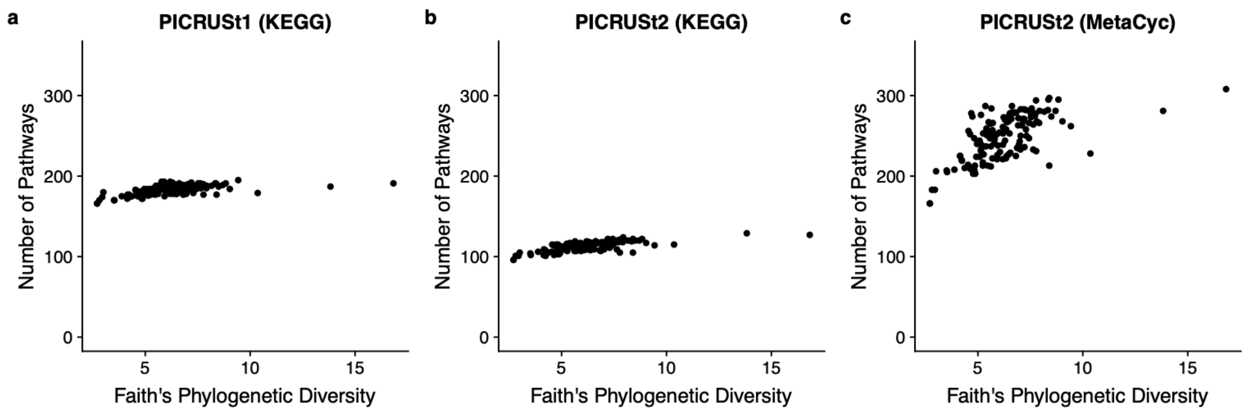


**Supplementary Figure 1**: The number of predicted pathways as phylogenetic diversity varies in samples from the Human Microbiome Project. A comparison of (a) KEGG pathways output by PICRUSt1, (b) KEGG pathways output by PICRUSt2, and (c) MetaCyc pathways output by the PICRUSt2 default pipeline. The number of KEGG pathways plateaus almost immediately whereas there is a much greater range in the MetaCyc pathways present. In addition, a mean of 1.6-fold more pathways are called as present in PICRUSt1 that are not called as present in PICRUSt2, which is due to the more stringent pathway pipeline intended to reduce false positives.

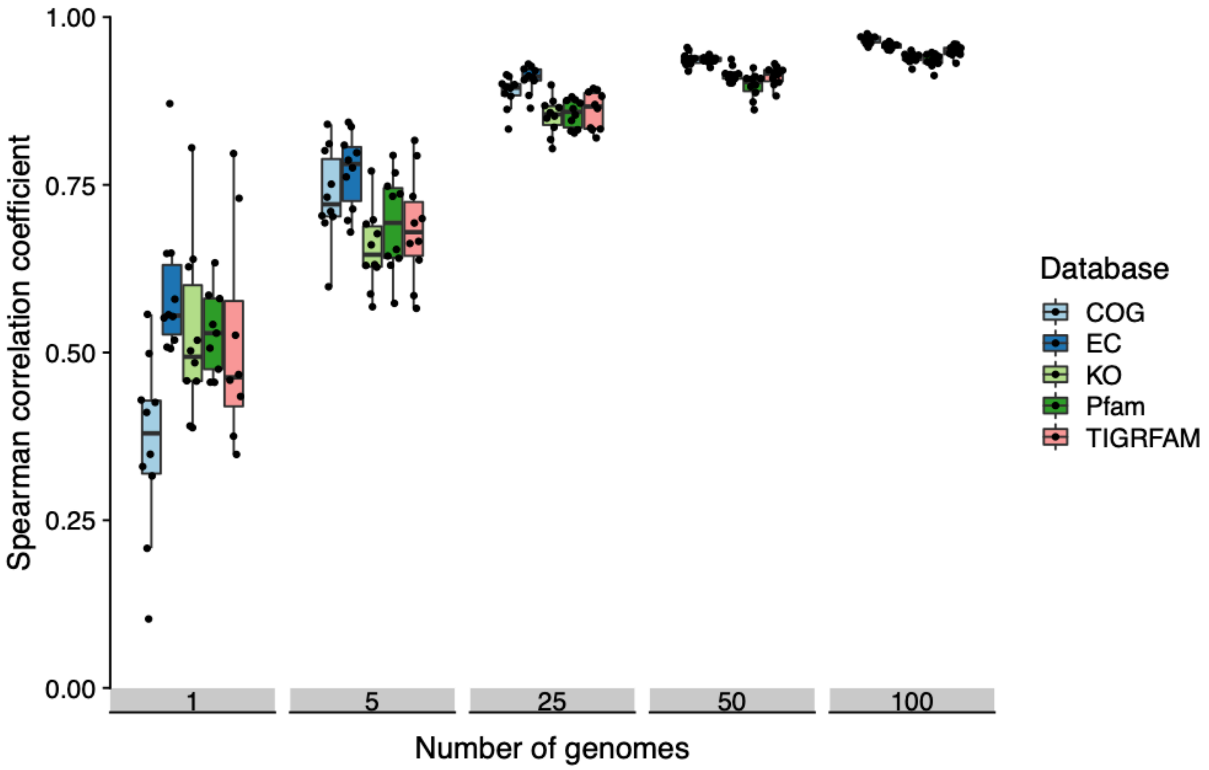


**Supplementary Figure 2**: Spearman correlations between number of gene families in two random subsets of genomes. As the size of the random subsets increases the correlation coefficient between the two subsets approaches 1. Ten replicates are plotted for each number of genome subsets. These correlations are shown for all functional databases that can be used with PICRUSt2 by default. Clusters of Orthologous Genes (COG); Enzyme Classification Numbers (EC); Kyoto Encyclopedia of Genes and Genomes Ortholog (KO); Protein Families (Pfam); and The Institute for Genomic Research's database of protein FAMilies (TIGRFAM).

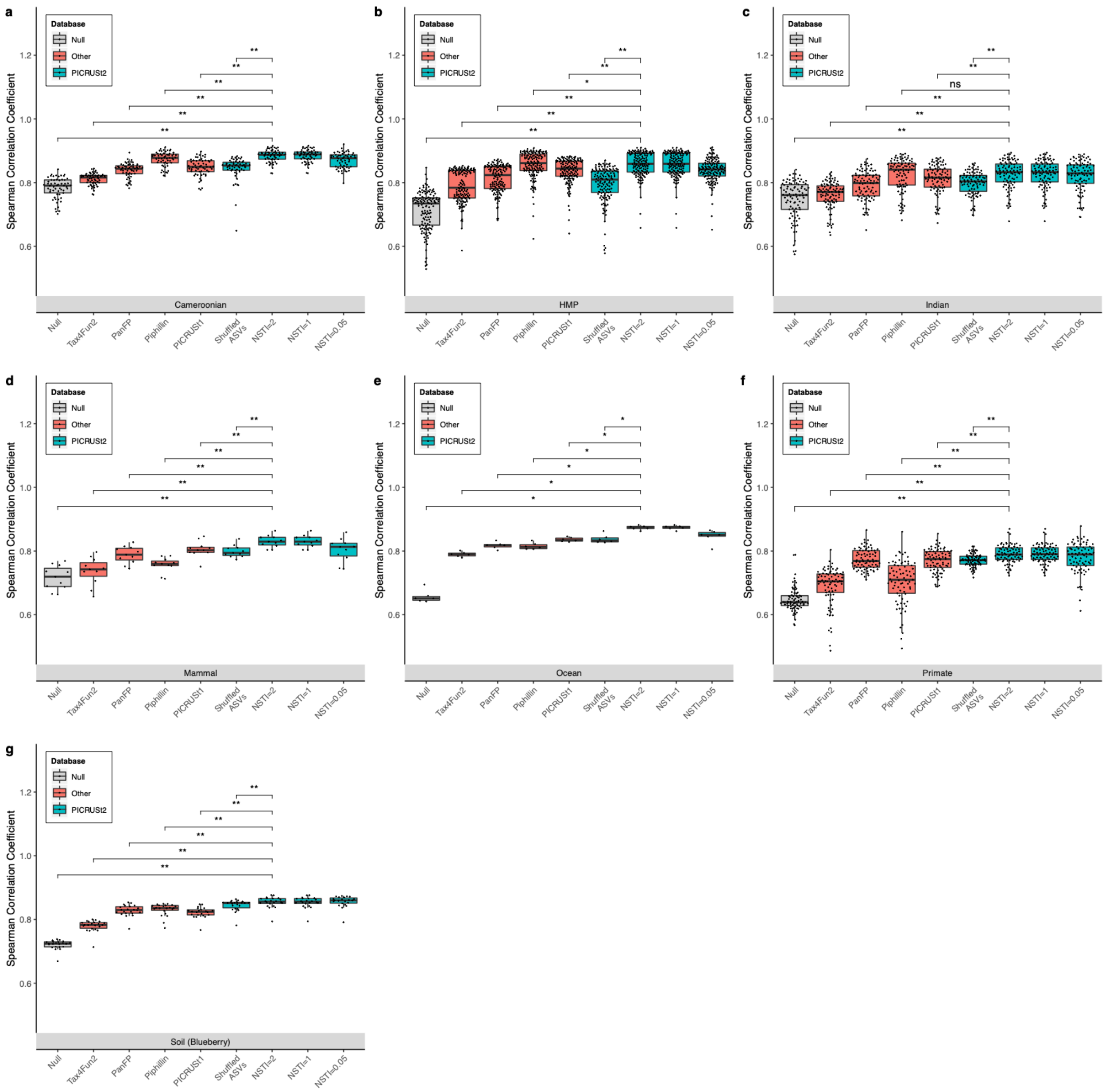


**Supplementary Figure 3:** Full KEGG ortholog validation results of PICRUSt2 comparing metagenome prediction performance against gold-standard shotgun metagenomic sequencing. The data shown here is the same as in Figure 2 except the performance at different nearest-sequenced taxon index (NSTI) cut-offs for the ASV input data is also shown. HMP: Human Microbiome Project. Significance of paired-sample, two-tailed Wilcoxon tests is indicated above each tested grouping (*, **, and ns correspond to $P < 0.05$, $P < 0.001$, and not significant respectively).

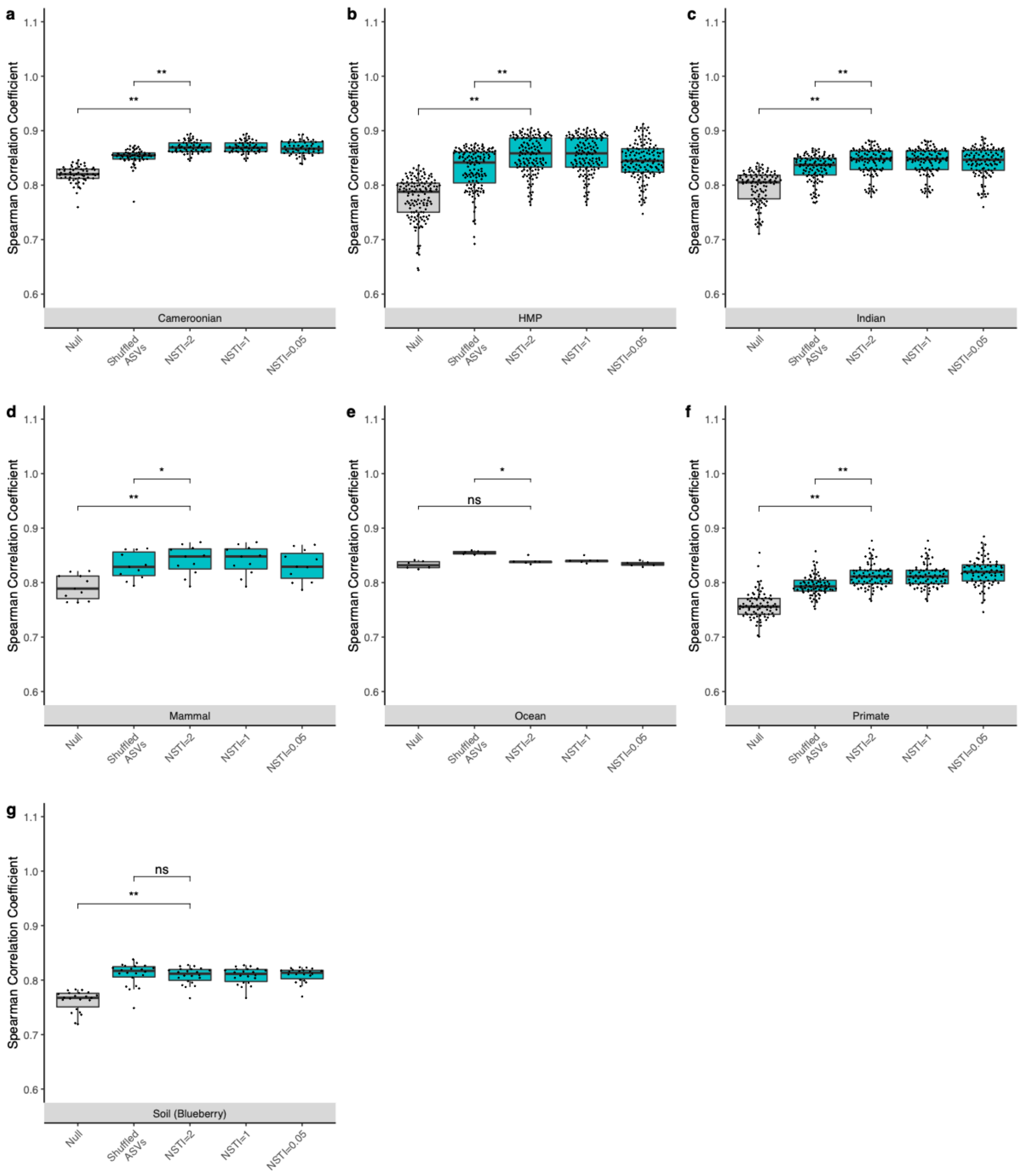


**Supplementary Figure 4:** Spearman correlation coefficients of predicted MetaCyc pathway abundances compared to shotgun metagenomics on same samples for (a) stool samples from Cameroonian individuals, (b) the human microbiome project (HMP), (c) stool samples from Indian individuals, (d) mammalian stool samples, (e) ocean samples, (f) non-human primate stool samples, and (g) soil samples. PICRUSt2 predictions based on varying nearest sequenced taxon index (NSTI) cut-offs are shown to illustrate how this parameter affects prediction performance. Significance of paired-sample, two-tailed Wilcoxon tests is indicated above each tested grouping (*, **, and ns correspond to $P < 0.05$, $P < 0.001$, and not significant, respectively).

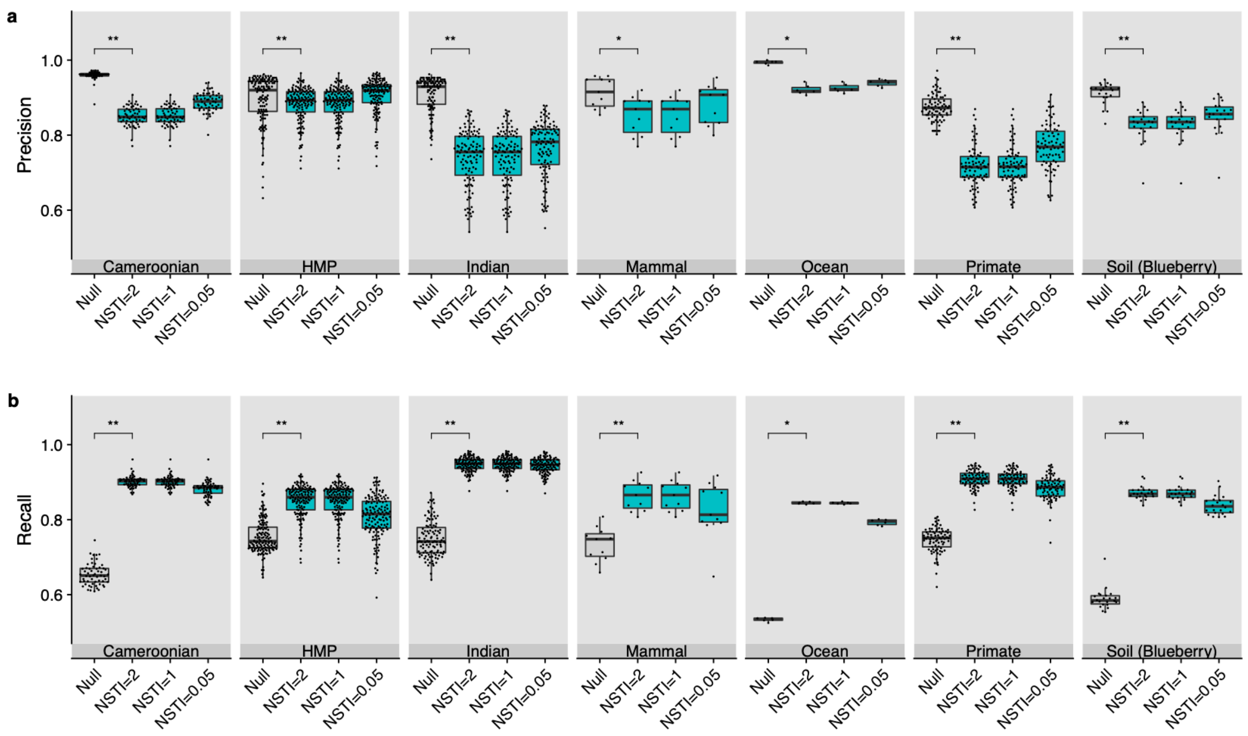


**Supplementary Figure 5:** (a) Precision and (b) recall of predicted MetaCyc pathway abundances compared to shotgun metagenomics on same samples for the Human Microbiome Project (HMP), mammalian stool samples, ocean samples, and soil samples. PICRUSt2 predictions based on varying nearest sequenced taxon index (NSTI) cut-offs are shown to illustrate how this parameter affects prediction performance. Significance of paired-sample, two-tailed Wilcoxon tests is indicated above each tested grouping (* and ** correspond to P < 0.05 and P < 0.001, respectively).

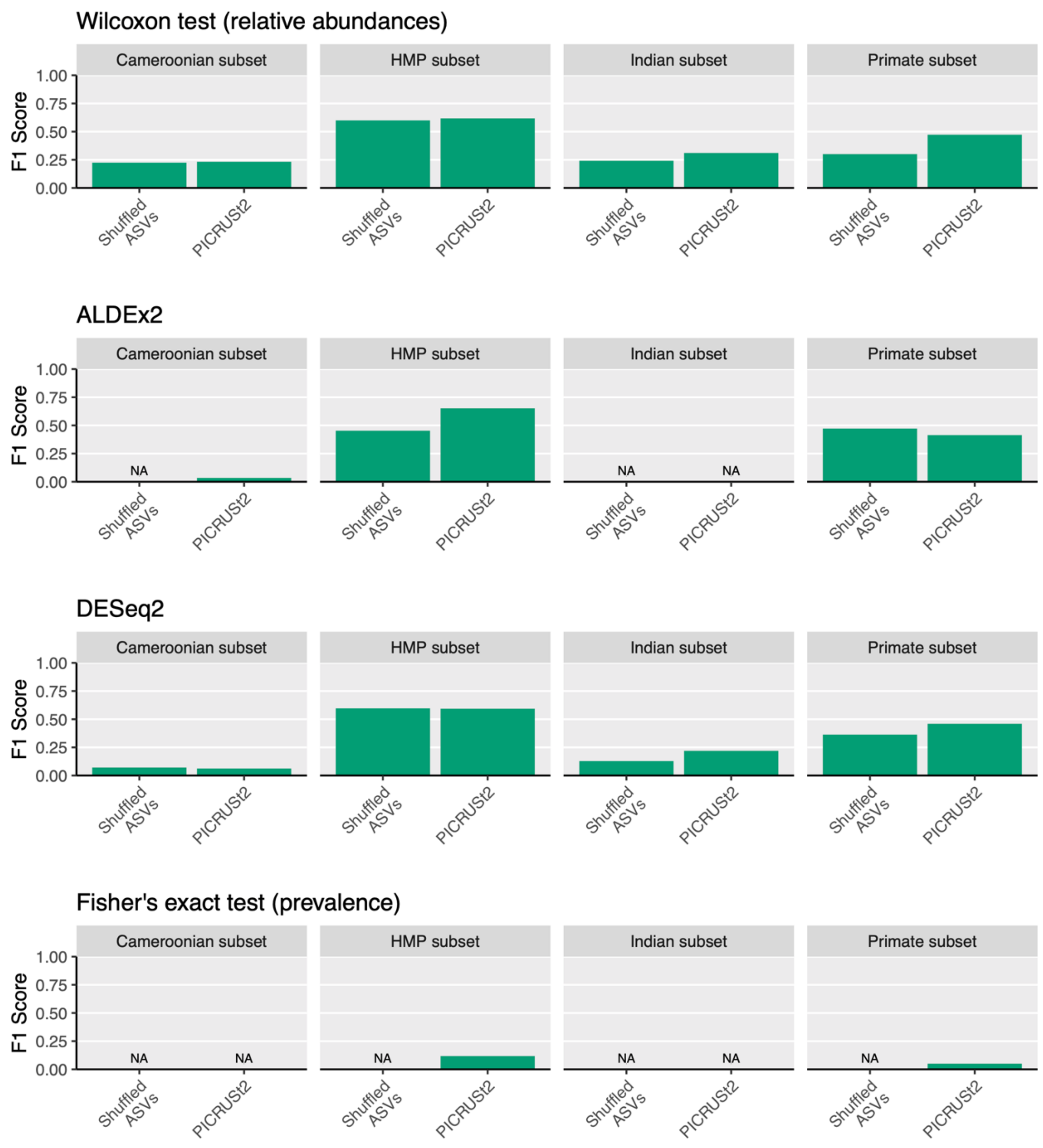


**Supplementary Figure 6:** Differential abundance and prevalence of predicted MetaCyc pathways vary across datasets and with statistical methods used and are similar to predictions based on shuffled ASVs. The F1 score is plotted to summarize the agreement between the statistical testing of predicted pathways with pathways identified from HUMAnN2. All pathway abundances for this analysis were based on Enzyme Classification numbers over the entire metagenome sample (i.e. assuming a bag-of-genes model), rather than restricting pathway predictions to each Amplicon Sequence Variant's (ASV) predicted genome. Each of these statistical tests has a different null hypothesis, but the first three methods are all commonly used to perform differential abundance analyses on microbiome data. NA indicates cases where either the recall or precision could not be calculated (which means the F1 scores was not applicable). The sample groupings for these analyses are the same as for all the other differential testing results reported and are indicated by "subset" here simply to save space.

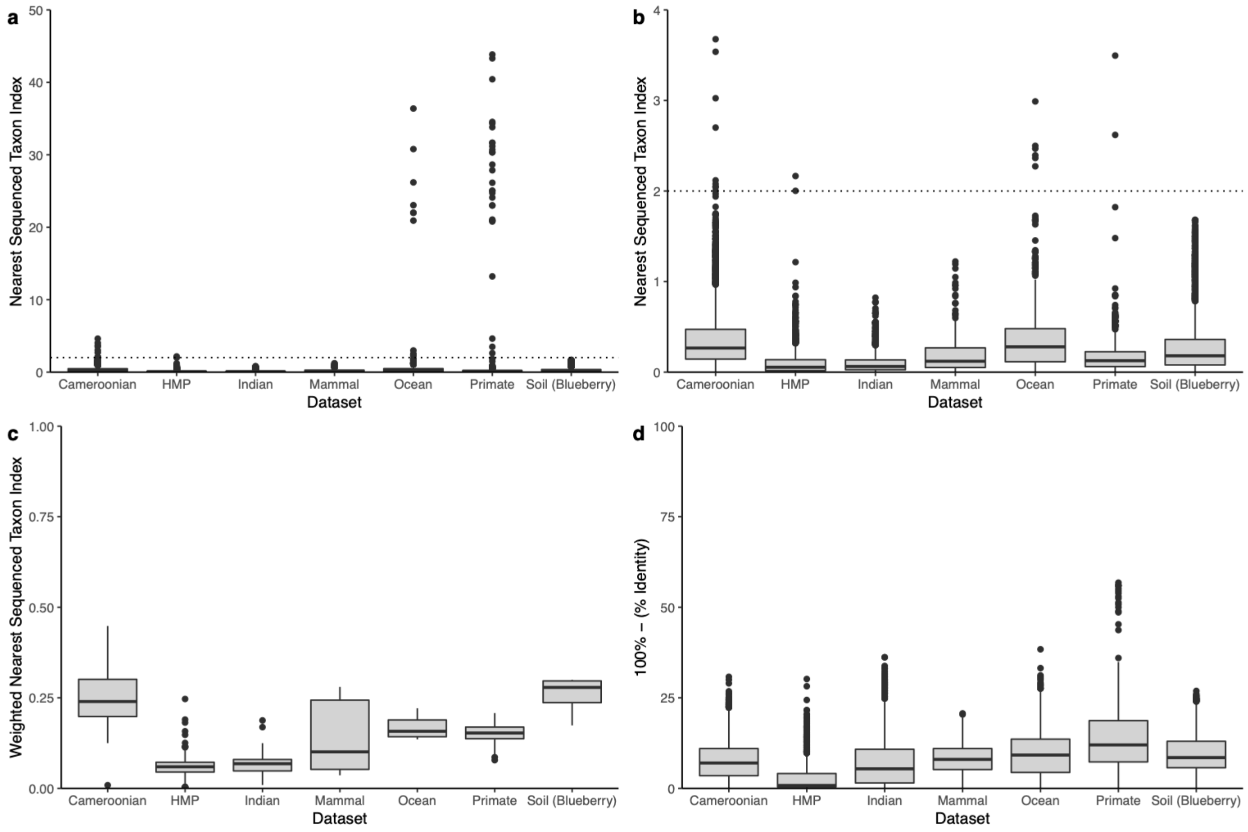


**Supplementary Figure 7**: Reference-based quality metrics for amplicon sequence variants (ASVs) of validation datasets. (a) Nearest sequenced taxon index (NSTI) for ASVs (branch length to nearest reference sequence). The grey horizontal dotted line at two indicates the default maximum NSTI value over which ASVs will be excluded. (b) Same as panel a except y-axis is truncated so visualizing these differences is easier. (c) Per-sample weighted NSTI values, which corresponds to the NSTI values in panels a and b under the maximum cut-off weighted by the abundance of each ASV. (d) Complement of percent identities of ASVs against reference database sequences.

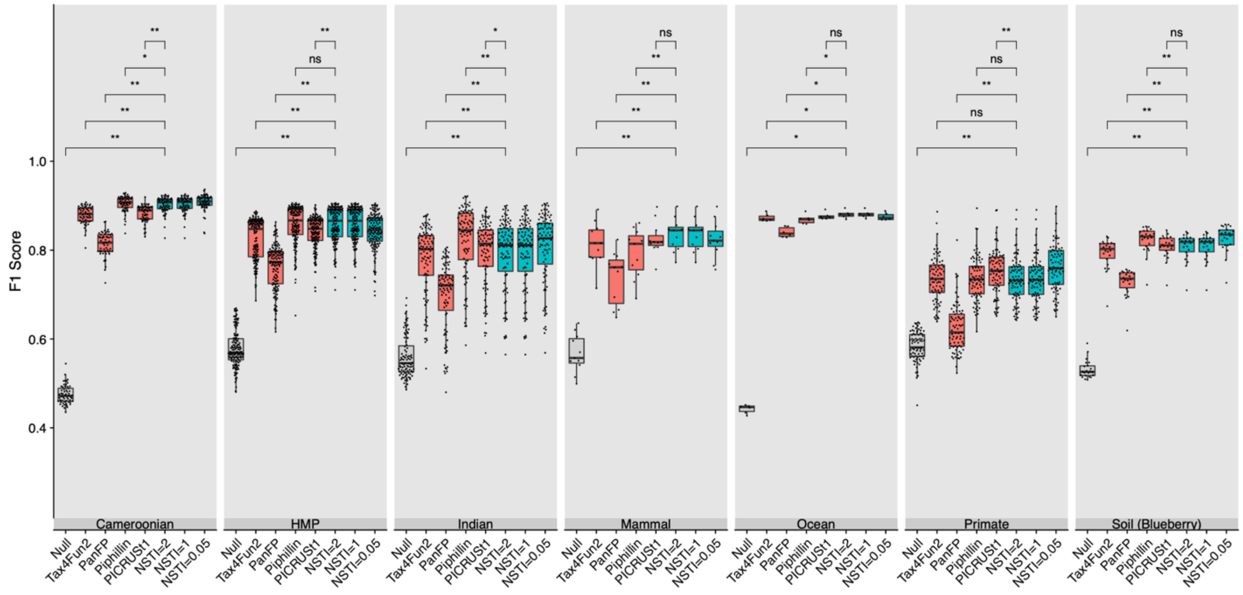


**Supplementary Figure 8**: KEGG ortholog prediction performance in terms of F1 score, which is the harmonic mean of precision and recall. The datasets and prediction categories shown here are the same as in Figure 2 except the PICRUSt2 performance at different nearest-sequenced taxon index (NSTI) cut-offs for the ASV input data is also shown. Boxplot colours refer to three different category groupings: the null expectation (grey), alternative prediction tools (red), and PICRUSt2 predictions (cyan). The significance of paired-sample, two-tailed Wilcoxon tests is indicated above each tested grouping (*, **, and ns correspond to $P < 0.05$, $P < 0.001$, and not significant respectively). HMP: Human Microbiome Project.

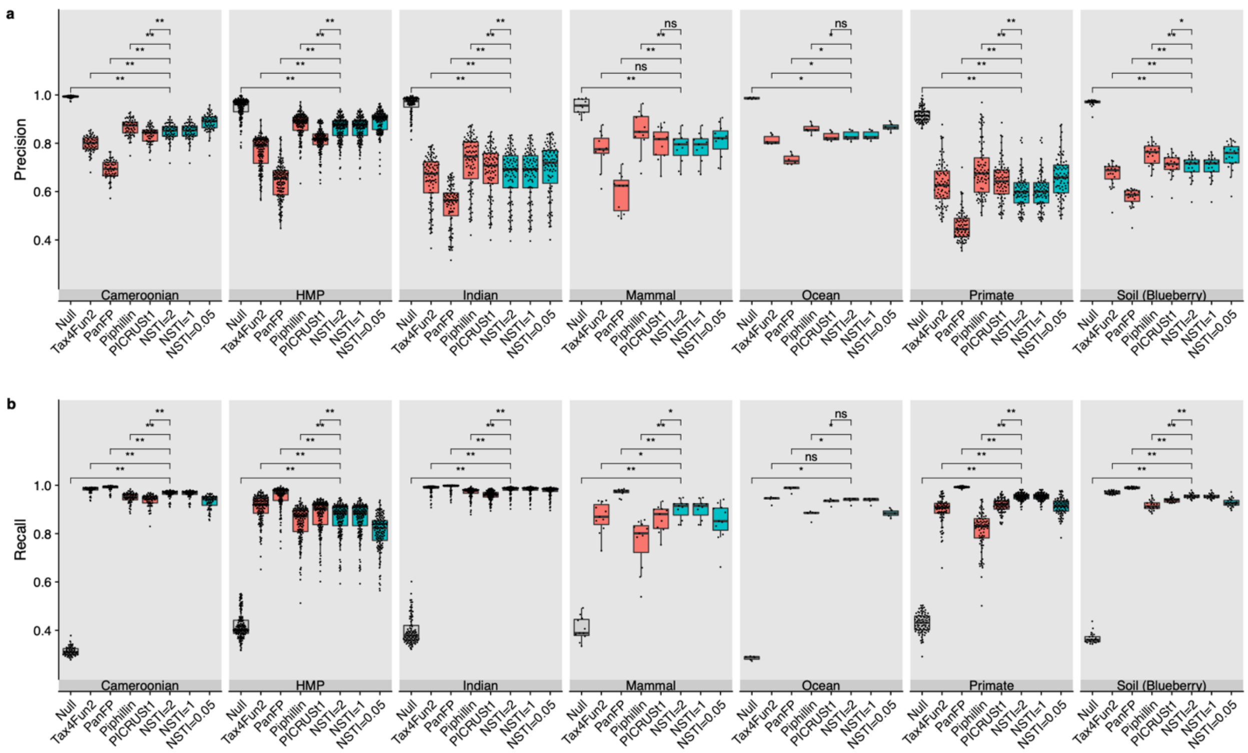


**Supplementary Figure 9**: KEGG ortholog prediction performance in terms of (a) precision and (b) recall. The datasets and prediction categories shown here are the same as in Figure 2 except the PICRUSt2 performance at different nearest-sequenced taxon index (NSTI) cut-offs for the ASV input data is also shown. Boxplot colours refer to three different category groupings: the null expectation (grey), alternative prediction tools (red), and PICRUSt2 predictions (cyan). The significance of paired-sample, two-tailed Wilcoxon tests is indicated above each tested grouping (*, **, and ns correspond to P < 0.05, P < 0.001, and not significant respectively). HMP: Human Microbiome Project.

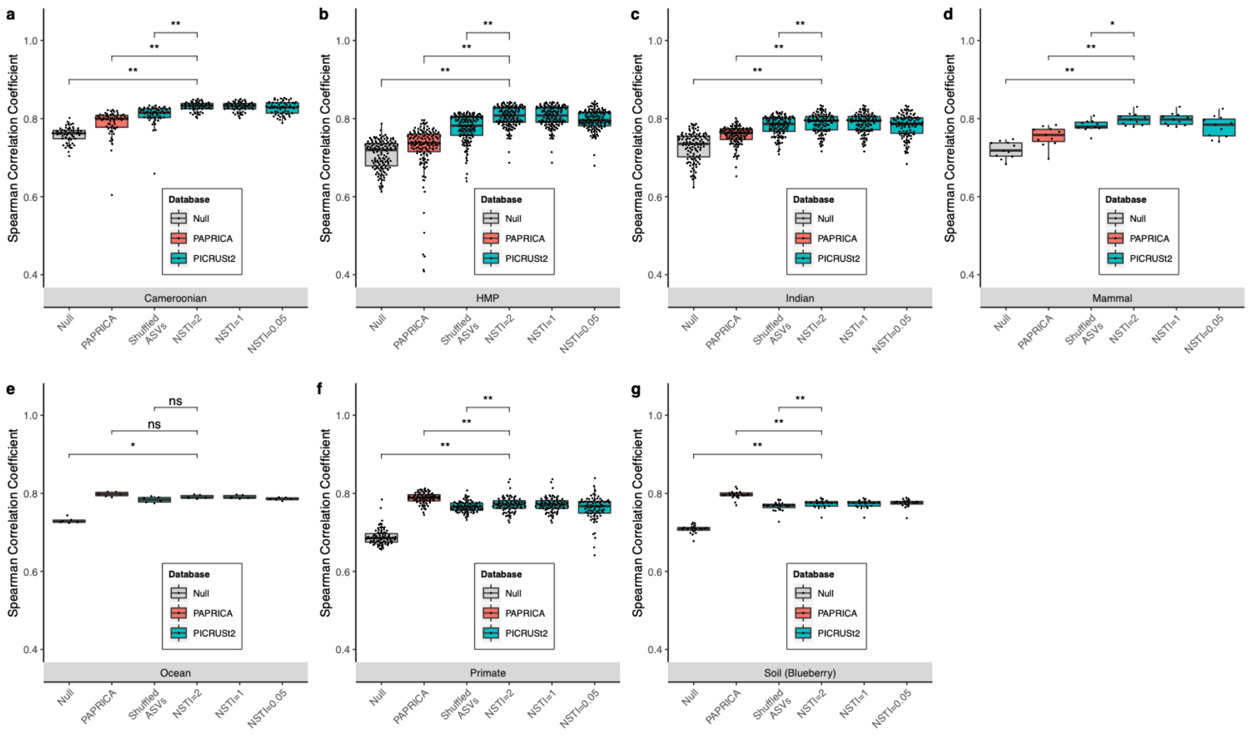


**Supplementary Figure 10:** Spearman correlation coefficients of predicted Enzyme Classification (EC) number abundances compared to shotgun metagenomics on same samples for (a) stool samples from Cameroonian individuals, (b) the human microbiome project (HMP), (c) stool samples from Indian individuals, (d) mammalian stool samples, (e) ocean samples, (f) non-human primate stool samples, and (g) soil samples. PICRUSt2 predictions based on varying nearest sequenced taxon index (NSTI) cut-offs are shown to illustrate how this parameter affects prediction performance. Significance of paired-sample, two-tailed Wilcoxon tests is indicated above each tested grouping (*, **, and ns correspond to $P < 0.05$, $P < 0.001$, and not significant, respectively).

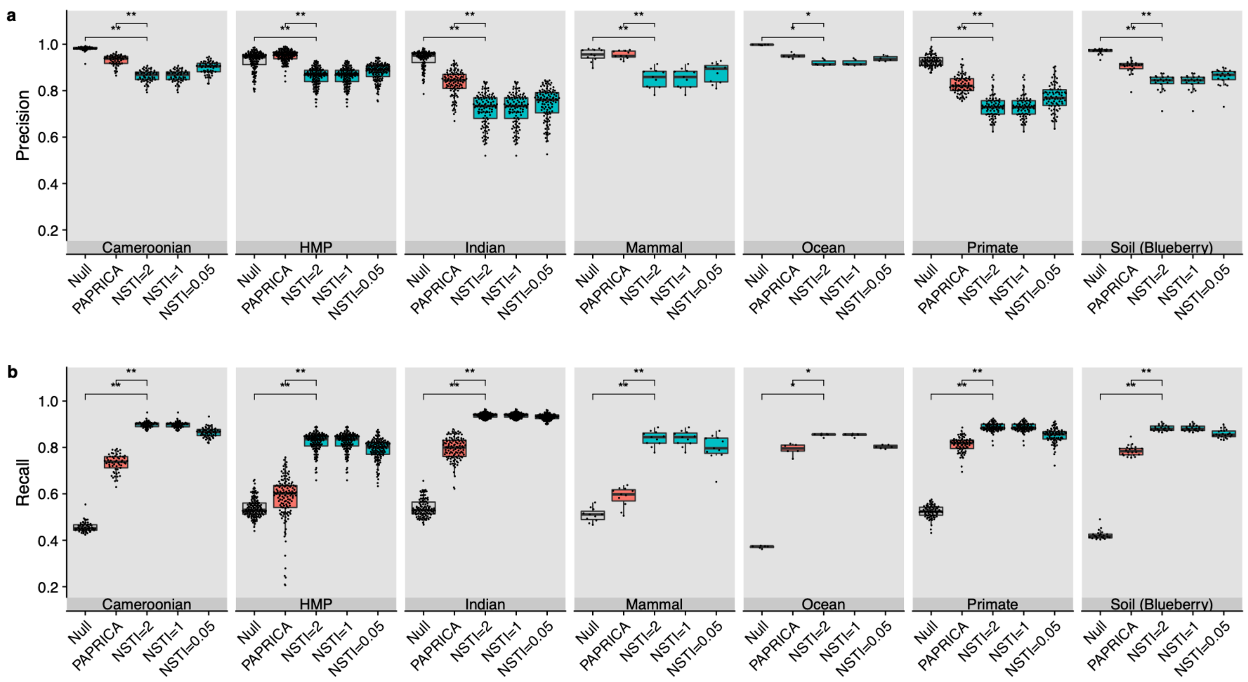


**Supplementary Figure 11:** (a) Precision and (b) recall of predicted Enzyme Classification (EC) numbers (based on presence/absence) compared to shotgun metagenomics for all seven validation datasets. PICRUSt2 predictions based on varying nearest sequenced taxon index (NSTI) cut-offs are shown to illustrate how this parameter affects prediction performance. Significance of paired-sample, two-tailed Wilcoxon tests is indicated above each tested grouping (* and ** correspond to $P < 0.05$ and $P < 0.001$, respectively).

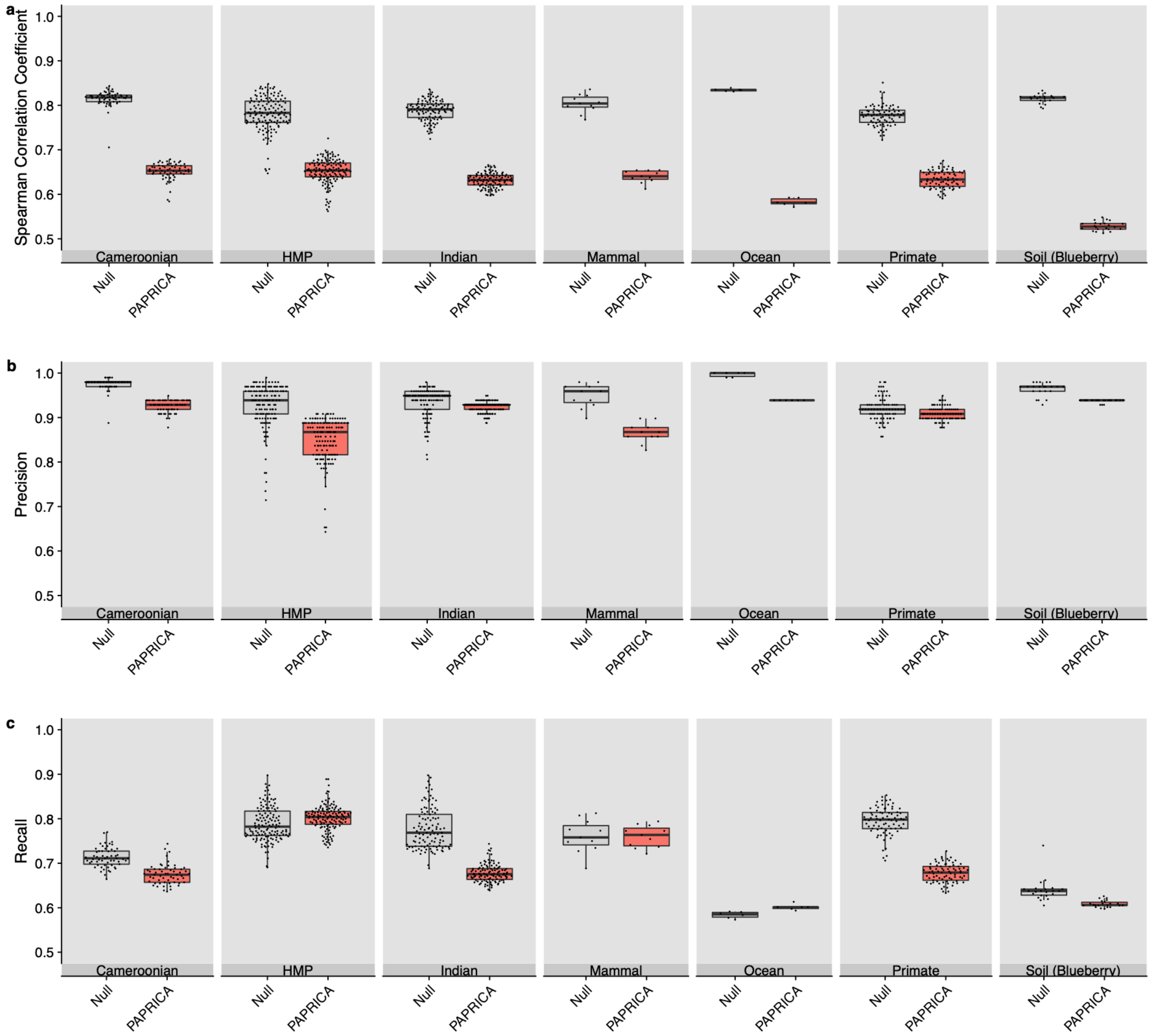


**Supplementary Figure 12**: Poor concordance of PAPRICA pathway abundance predictions compared to pathways identified by HUMAnN2 in shotgun metagenomics data based on (a) Spearman correlation coefficients, (b) precision, and (c) recall. Only 193 MetaCyc pathways are considered here since this is the only set that could be identified by both PAPRICA and HUMAnN2. In contrast, 575 pathways could potentially be identified by both PICRUSt2 and HUMAnN2. The poor concordance shown here may be due to the differing approach for inferring pathway levels used by PAPRICA, which may mean that comparing these predictions with HUMAnN2 is not a fair evaluation.

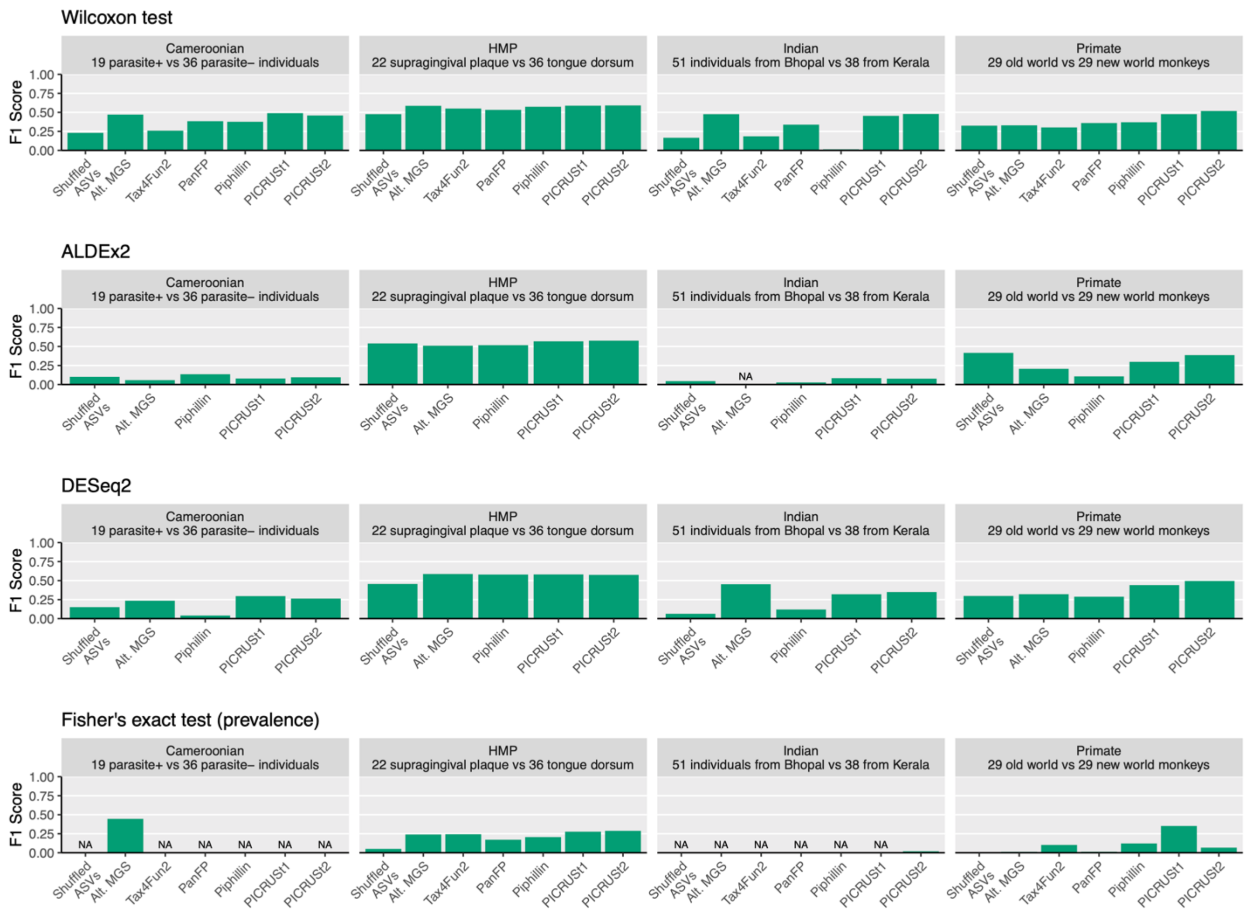


**Supplementary Figure 13:** Differential abundance and prevalence of PICRUSt2 predicted KEGG orthologs (KOs) agree only marginally, but best overall, with shotgun metagenomics data. The F1 score is plotted to summarize the agreement between the statistical testing of predicted KOs with KOs identified from HUMAnN2. PanFP and Tax4Fun2 are missing from the ALDEx2 and DESeq2 results because these statistical methods require count tables as input. Each of these statistical tests has a different null hypothesis, but the first three methods are all commonly used to perform differential abundance analyses on microbiome data. NA indicates cases where either the recall or precision could not be calculated (which means the F1 scores was not applicable). “Alt. MGS” refers to the alternative shotgun metagenomics processing pipeline.

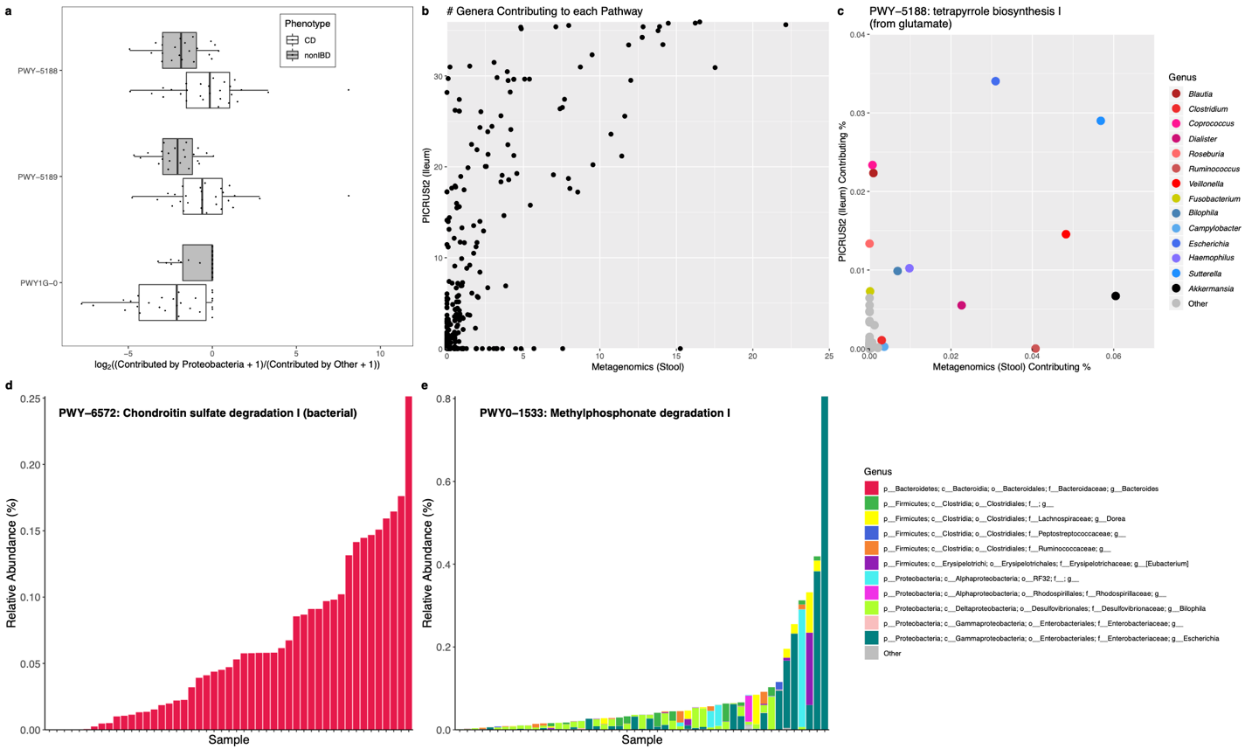


**Supplementary Figure 14:** Applying PICRUSt2 to Crohn's disease cohort yields novel insights. (a) Predicted MetaCyc pathways significantly contributed by Proteobacteria in the ileum of subjects with Crohn's disease (CD, n=27) compared to healthy controls (non-IBD, n=20) based on PICRUSt2 inference from 16S rRNA gene sequencing. These significant pathways are: PWY-5188 (tetrapyrrole biosynthesis I [from glutamate]), PWY-5189 (tetrapyrrole biosynthesis II [from glycine]), and PWY1G-0 (mycothiol biosynthesis). (b) The mean number of classified genera contributing to each of the 313 MetaCyc pathways identified in either the ileum 16S rRNA gene sequencing (by PICRUSt2) or shotgun metagenomics sequencing (MGS) of the stool of the same CD subjects. (c) The top classified genera contributing to the relative abundance of PWY-5188 based on PICRUSt2 predictions of 16S rRNA gene sequencing in ileum tissue or MGS of the stool of the same subjects. Only the top 10 contributing genera are shown. Genera of the same phylum are shades of the same colour (Firmicutes and Proteobacteria are shades of red and blue, respectively). (d and e) Taxonomic breakdown of genera contributing to the predicted relative abundance of (d) PWY-6572 and (e) PWY0-1533 across all CD samples. Unclassified genera were included in these stacked bar charts, unlike in panel c. Only the top ten genera contributing to either PWY-6572 or PWY0-1533 are labelled.

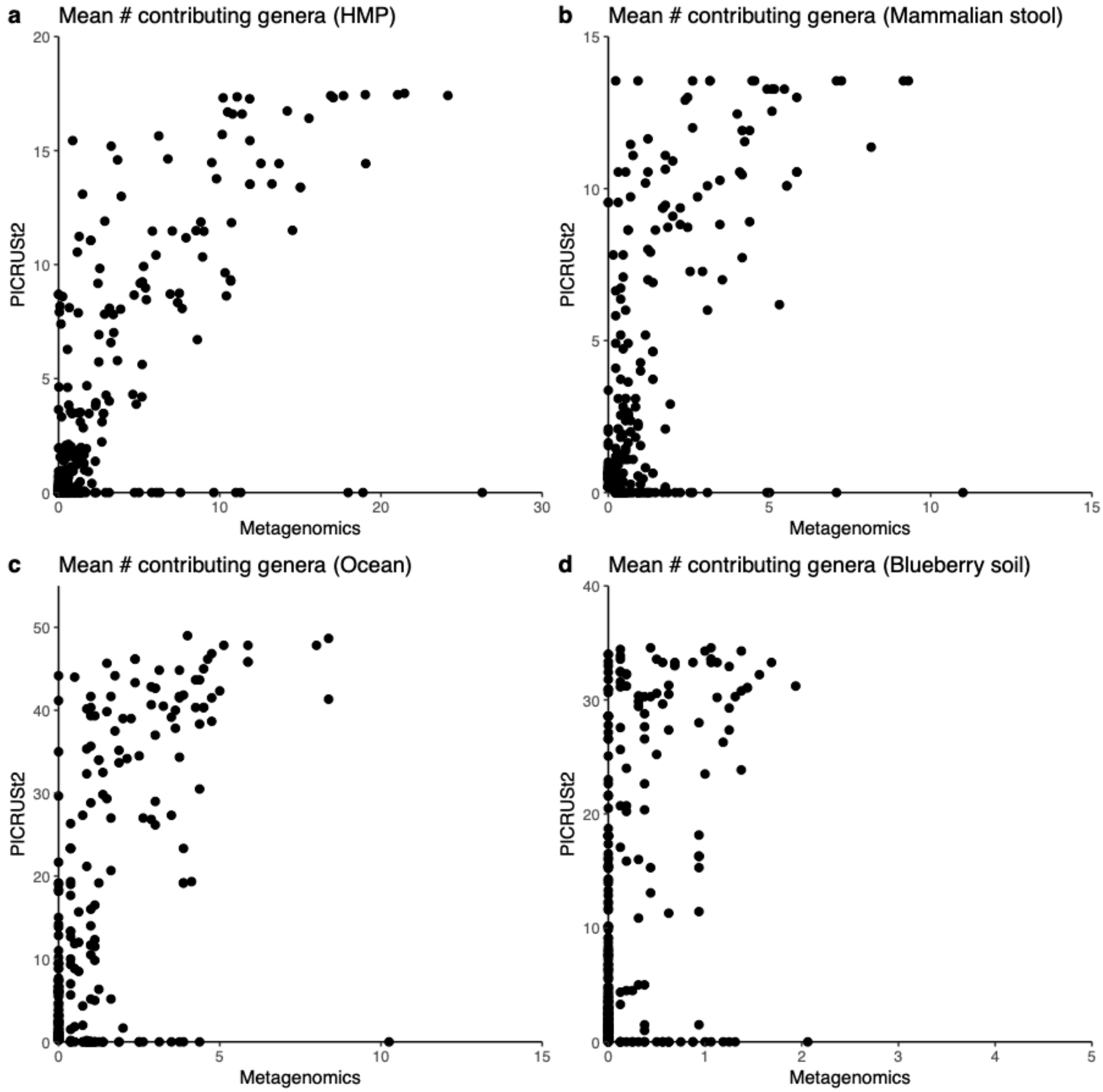


**Supplementary Figure 15:** The mean number of classified genera contributing for every MetaCyc pathway identified in either the 16S rRNA gene sequencing (by PICRUSt2) or shotgun metagenomics sequencing (MGS) of four representative datasets: (a) the Human Microbiome Project (HMP), (b) mammalian stool, (c) ocean, or (d) blueberry soil samples. This figure is meant to complement the analysis performed on the Crohn's disease biopsy data, because in this case the comparison is made between 16S rRNA gene sequencing and MGS on the same samples. Note that the number of characterized genera could be affected by sampling environment as labelled here, but importantly these datasets also differ in several technological ways as outlined in Supplementary Table 2.

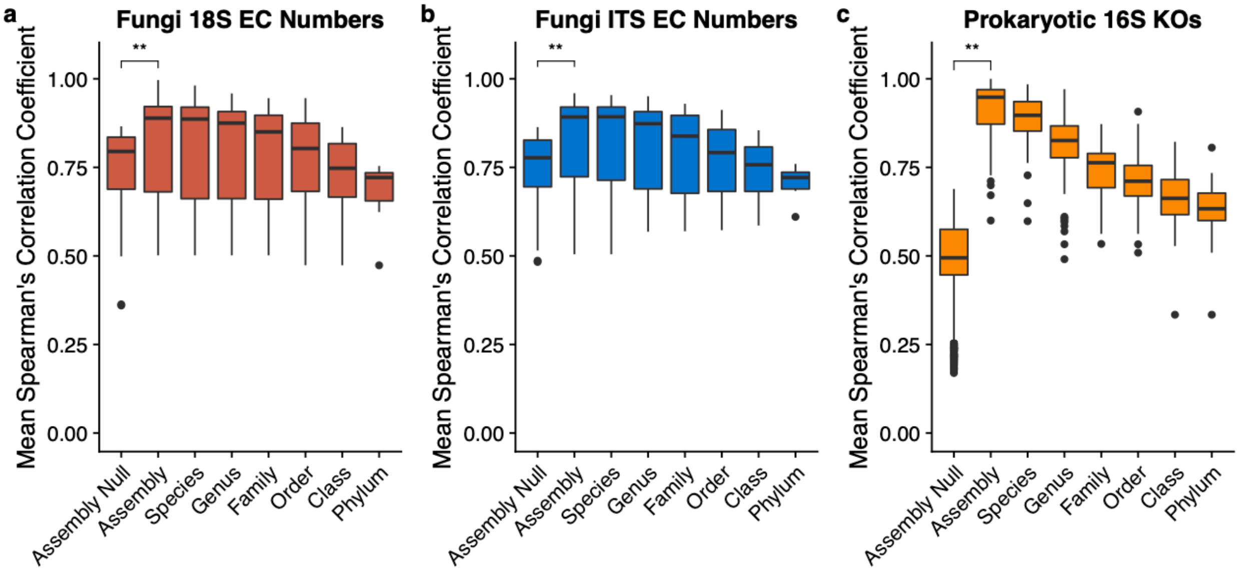


**Supplementary Figure 16:** Validating test fungi databases and key prokaryotic database discussed in main-text with genome holdout analysis: (A) fungi 18S rRNA gene-predicted Enzyme Classification (EC) Numbers, fungi internal transcribed spacer (ITS)-predicted EC Numbers, and (C) 16S rRNA gene-predicted KEGG orthologs (KOs). The prokaryotic database is included here as a point of reference for the fungi databases. For each database shown above, predictions were made for all genomes within each clade at a given taxonomic level after pruning all those genomes from the reference tree. The mean spearman correlation coefficient between the predicted and expected gene family abundances was then calculated for each clade. The "Assembly" level refers to individual genomes. The "Assembly Null" category corresponds to the correlation between the gene family abundances for each genome and the mean abundance of gene families across all genomes. The ** annotation indicates a significant Wilcoxon test ($P < 0.001$). Note that for the prokaryotic KO analysis that a maximum of 100 clades at each taxonomic level were selected randomly for this analysis to decrease computation time. In this plot the points correspond to outliers outside the boxplot whiskers only and all other points are not shown.

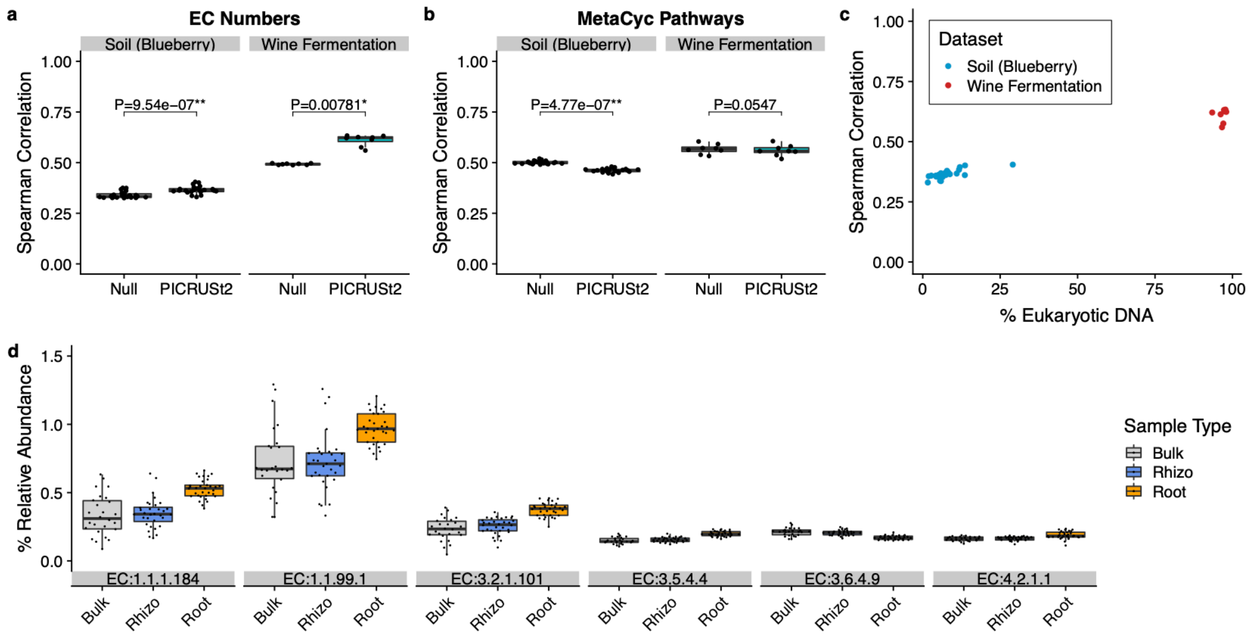


**Supplementary Figure 17:** PICRUSt2 18S rRNA gene and internal transcribed spacer predictions exceed null prediction accuracy. (A) Spearman correlation coefficients between amplicon predicted Enzyme Classification number abundances and gold-standard shotgun metagenomic (MGS) profiles from the same biological samples. (B) Spearman correlation coefficients between amplicon-predicted MetaCyc pathway abundances and MGS on the same biological samples. For panels A and B, the P-values of paired-sample, two-tailed Wilcoxon tests is indicated above each tested grouping (* and ** correspond to P < 0.05 and P < 0.001, respectively). (C) The Spearman correlation coefficients as shown in panel A re-plotted against the percent of non-animal and non-plant eukaryotic DNA within each sample. The blueberry soil dataset consists of 22 18S rRNA gene sequencing samples and the wine fermentation dataset consists of eight internal transcribed spacer region one (ITS1) sequencing samples. (D) The relative abundance of significantly informative EC numbers (P < 0.001) in a Random Forest model for distinguishing blueberry soil and root samples by sample type. Only significant EC numbers with a mean relative abundance greater than 0.15% are shown. The EC numbers shown correspond to carbonyl reductase (NADPH) (EC:1.1.1.184), choline dehydrogenase (EC:1.1.99.1), mannan endo-1,6-alpha-mannosidase (EC:3.2.1.101), adenosine deaminase (EC:3.5.4.4), chaperonin ATPase (EC:3.6.4.9), and carbonate dehydratase (EC:4.2.1.1). There are 26, 33, and 32 samples for the bulk, rhizosphere (rhizo), and root environments, respectively.

**Supplementary Table 1: Summary statistics of PICRUSt2 trait reference databases**

| Database | # Categories | Mean | sd | Sparsity | Mean trait depth |
|---|---|---|---|---|---|
| 16S rRNA gene | 1 | 2.51 | 2.45 | 0.00 | 2.52 |
| COG | 4598 | 0.55 | 1.76 | 0.68 | 0.54 |
| TIGRFAM | 4287 | 0.31 | 0.98 | 0.77 | 0.48 |
| EC | 2913 | 0.33 | 0.95 | 0.78 | 0.43 |
| PFAM | 11089 | 0.40 | 2.10 | 0.82 | 0.40 |
| KEGG | 10543 | 0.19 | 0.65 | 0.85 | 0.32 |

**Supplementary Table 2: Descriptions of the four paired 16S rRNA gene and shotgun metagenomics sequencing validation datasets used**

| Dataset | N* | 16S pipeline | 16S region | 16S tech. | MGS tech. | # ASVs | Mean # 16S reads | Mean # MGS reads | 16S read length | MGS read length |
|---|---|---|---|---|---|---|---|---|---|---|
| Cameroon | 57 | Deblur | V5-V6 | Illumina MiSeq | Illumina HiSeq | 4077 | 95354 | $42 \times 10^6$ | 228 | 101 |
| India | 91 | Deblur | V3 | Illumina NextSeq 500 | Illumina NextSeq 500 | 2237 | 164847 | $6.5 \times 10^6$ | 130 | 151 |
| HMP | 137 | DADA2 | V4 | Roche 454 | Illumina HiSeq | 1576 | 4592 | $30.0 \times 10^6$ | 245 | 101 |
| Primate | 77 | Deblur (QIITA) | V4 | Illumina MiSeq | Illumina HiSeq | 7452 | 15536 | $7.5 \times 10^6$ | 150 | 160 |
| Mammal | 8 | Deblur | V6-V8 | Illumina MiSeq | Illumina MiSeq | 323 | 2906 | $8.0 \times 10^6$ | 400 | 151 |
| Ocean | 6 | Deblur | V4 | Illumina MiSeq | Illumina HiSeq | 1148 | 62498 | $64.1 \times 10^6$ | 250 | 101 |
| Soil | 22 | Deblur | V6-V8 | Illumina MiSeq | Illumina NextSeq | 3333 | 5550 | $26.4 \times 10^6$ | 404 | 150 |

*Final number of samples overlapping between 16S rRNA gene and MGS datasets used for analyses.
Note: ASV and read descriptions correspond to final filtered data, “16S” refers to 16S rRNA gene, and “MGS” refers to shotgun metagenomics.

**Supplementary Table 3: Predicted pathways significantly associated (based on partial Spearman correlation [R]) with gene expression levels of Crohn's disease biomarkers in ileal tissue of subjects with Crohn's disease**

| Pathway | Gene | R | p | FDR |
|---|---|---|---|---|
| CALVIN-PWY | MMP3 | 0.6134 | 0.0009 | 0.0563 |
| GLUCOSE1PMETAB-PWY | MMP3 | -0.6024 | 0.0011 | 0.0677 |
| GLYOXYLATE-BYPASS | DUOX2 | -0.6018 | 0.0011 | 0.0677 |
| GLYOXYLATE-BYPASS | MMP3 | -0.7074 | 0.0001 | 0.0330 |
| HEME-BIOSYNTHESIS-II | DUOX2 | -0.6534 | 0.0003 | 0.0545 |
| HEME-BIOSYNTHESIS-II | MMP3 | -0.6214 | 0.0007 | 0.0545 |
| HEMESYN2-PWY | MMP3 | -0.6168 | 0.0008 | 0.0545 |
| PWY0-1241 | DUOX2 | -0.5757 | 0.0021 | 0.0956 |
| PWY0-1261 | MMP3 | -0.5799 | 0.0019 | 0.0945 |
| PWY0-1533 | MMP3 | -0.6203 | 0.0007 | 0.0545 |
| PWY-5097 | DUOX2 | 0.7074 | 0.0001 | 0.0330 |
| PWY-5154 | MMP3 | -0.6278 | 0.0006 | 0.0545 |
| PWY-5189 | DUOX2 | -0.5876 | 0.0016 | 0.0862 |
| PWY-5855 | DUOX2 | -0.6169 | 0.0008 | 0.0545 |
| PWY-5855 | MMP3 | -0.6183 | 0.0008 | 0.0545 |
| PWY-5856 | DUOX2 | -0.6169 | 0.0008 | 0.0545 |
| PWY-5856 | MMP3 | -0.6183 | 0.0008 | 0.0545 |
| PWY-5857 | DUOX2 | -0.6169 | 0.0008 | 0.0545 |
| PWY-5857 | MMP3 | -0.6183 | 0.0008 | 0.0545 |
| PWY-5918 | DUOX2 | -0.6203 | 0.0007 | 0.0545 |
| PWY-5918 | MMP3 | -0.5766 | 0.0020 | 0.0956 |
| PWY-6572 | NAT8 | -0.5743 | 0.0022 | 0.0956 |
| PWY-6708 | DUOX2 | -0.6169 | 0.0008 | 0.0545 |
| PWY-6708 | MMP3 | -0.6183 | 0.0008 | 0.0545 |
| PWY-6737 | DUOX2 | 0.6500 | 0.0003 | 0.0545 |
| PWY-6737 | MMP3 | 0.6446 | 0.0004 | 0.0545 |
| PWY-7184 | MMP3 | -0.5709 | 0.0023 | 0.0994 |
| PWY-7234 | MMP3 | -0.5931 | 0.0014 | 0.0794 |
| SO4ASSIM-PWY | MMP3 | -0.5852 | 0.0017 | 0.0873 |

**Supplementary Table 4: Counts of taxa in the tested internal transcribed spacer (ITS) and 18S rRNA gene databases**

| Database | Phyla | Classes | Orders | Families | Genera | Species | Genomes |
|---|---|---|---|---|---|---|---|
| ITS | 7 | 26 | 48 | 89 | 130 | 174 | 183 |
| 18S rRNA gene | 8 | 27 | 52 | 102 | 151 | 193 | 209 |
